# Categorical edge-based analyses of phylogenomic data reveal conflicting signals for difficult relationships in the avian tree

**DOI:** 10.1101/2021.05.17.444565

**Authors:** Ning Wang, Edward L. Braun, Bin Liang, Joel Cracraft, Stephen A. Smith

## Abstract

Phylogenetic analyses fail to yield a satisfactory resolution of some relationships in the tree of life even with genome-scale datasets, so the failure is unlikely to reflect limitations in the amount of data. Gene tree conflicts are particularly notable in studies focused on these contentious nodes, and taxon sampling, different analytical methods, and/or data type effects can further confound analyses. Although many efforts have been made to incorporate biological conflicts, few studies have curated individual genes for their efficiency in phylogenomic studies. Here, we conduct an edge-based analysis of Neoavian evolution, examining the phylogenetic efficacy of two recent phylogenomic bird datasets and three datatypes (ultraconserved elements [UCEs], introns, and coding regions). We assess the potential causes for biases in signal-resolution for three difficult nodes: the earliest divergence of Neoaves, the position of the enigmatic Hoatzin *(Opisthocomus hoazin),* and the position of owls (Strigiformes). We observed extensive conflict among genes for all data types and datasets even after meticulous curation. Edge-based analyses (EBA) increased congruence and provided information about the impact of data type, GC content variation (GC_CV_), and outlier genes on each of nodes we examined. First, outlier gene signals appeared to drive different patterns of support for the relationships among the earliest diverging Neoaves. Second, the placement of Hoatzin was highly variable, although our EBA did reveal a previously unappreciated data type effect with an impact on its position. It also revealed that the resolution with the most support here was Hoatzin + shorebirds. Finally, GCCV, rather than data type (i.e., coding vs non-coding)*per se,* was correlated with a signal that supports monophyly of owls + Accipitriformes (hawks, eagles, and New World vultures). Eliminating high GC_CV_ loci increased the signal for owls + mousebirds. Categorical EBA was able to reveal the nature of each edge and provide a way to highlight especially problematic branches that warrant a further examination. The current study increases our understanding about the contentious parts of the avian tree, which show even greater conflicts than appreciated previously.

## 1. Introduction

The growth of genomic datasets is increasingly facilitating the resolution of phylogenetic relationships, although as datasets have grown, so has conflict. This may involve gene-tree discordances within a single study (e.g., Wang et al. 2019) or the resolution of difficult and controversial hypotheses found across multiple studies (e.g., Wickett et al. 2014; Puttick et al. 2018). Conflicting gene trees can be the result of biological processes such as hybridization, gene duplication and loss, and incomplete lineage sorting (ILS; see Table 1 for list of abbreviations) or they can be the result of artificial noise arising from, for example, assembly error, orthology inference, sequence alignment, model specification, and insufficient informative variation (Springer and Gatesy 2019). In part, due to these conflicts, several areas across the tree of life have consistently been difficult to resolve (Tarver et al. 2016; Shen et al. 2017) despite the use of genome-scale datasets (Foley et al. 2016) and increased taxa sampling (Prum et al. 2015; Wang et al. 2017) that benefit from growing effort and worldwide collaboration, as well as easier acquirement of ancient DNA (e.g., Chen et al. 2018).

**Table 1.** Glossary of terms and abbreviations used in this study.

| Term | Full name and explanation |
| --- | --- |
| EBA | edge-based analyses: constraint phylogenetic analyses on certain edge on the tree. |
| categorical EBA | categorical edge-based analyses |
| UCEs | Ultra-conserved elements |
| GCcv | GC content variation |
| ILS | Incomplete Lineage Sorting |
| BS | Bootstrap Support |
| UFBS | Ultra Fast Bootstrap Support |
| SH-aLRT | approximate likelihood-ratio test with the nonparametric SH correction |
| NNI | Nearest-neighbor interchange, which exchanges the connectivity of four subtrees within the main tree. |
| RF | Robinson-Foulds distances |
| TENT ExaML | the concatenated total evidence nucleotide tree from Jarvis et al. (2014) |
| ICA value | Internode Certainty All; the degree of certainty for a bipartition in a given set of trees in conjunction with all conflicting bipartitions in the same tree set |
| GTEE | gene-tree estimation error |
| $\ln L$ | log likelihood score |
| $\Delta \ln L$ | difference in $\ln L$ between the best and the second-best resolutions for each gene |
| $\sum \Delta \ln L$ | summation of $\Delta \ln L$ for each constraint from supporting genes |
| $\text{sig} \Delta \ln L$ | genes showing significant support for a specific topology (i.e., $\Delta \ln L > 2$) |
| $\sum \text{sig} \Delta \ln L$ | summation of $\Delta \ln L$ among genes with significant support (i.e., $\Delta \ln L > 2$) |
| $\sum N_g$ | the number of genes supporting a specific topology |
| EL/EH | exons with lower or higher $\Delta \text{GC}_{90}$/tree length |
| IL/IH | introns with lower or higher $\Delta \text{GC}_{90}$/tree length |
| UL/UH | UCEs with lower or higher $\Delta \text{GC}_{90}$/tree length |
| $\Delta \text{GC}_{50}$ | the GC differences between taxa with high (the upper quartile) and low (the lower quartile) GC content. |
| $\Delta \text{GC}_{90}$ | the GC differences between taxa with high (the 95 percentile) and low (the 5 percentile) GC content. |
| $\Delta \text{GC}_{100}$ | the GC differences between taxa with the highest and the lowest GC content. |
| Avian lineages: | Also see Jarvis et al. (2014) and Sangster et al. (2022) for clade names |
| *S* | Shorebirds (Charadriiformes) |
| *CR* | Cranes and Rails (Gruiformes) |
| *TB* | Turacos and Bustards (Musophagotides) |
| *M* | Mousebirds (Coliiformes) |
| *W* | Woodpeckers and allies (Cavitaves) |
| *EHV* | Eagles, Hawks, and New World Vultures (Accipitriformes) |
| “Magnificent Seven” | Superordinal groups in Neoaves tree that have been recovered in almost all recent large-scale studies; we use Roman numerals to indicate these groups |
| **I** | Telluraves (Core Landbirds) |
| **II** | Aequornithes (Core Waterbirds) |
| **III** | Phaethontimorphae (Sunbittern, Kagu, and Tropicbirds) |
| **IV** | Otidimorphae (Cuckoos, Bustards, and Turacos) |
| **V** | Strisores/Caprimulgimorphae (Nightjars, Swifts, Hummingbirds, and allies) |
| **VI** | Columbimorphae (Doves, Mesites, and Sandgrouse) |
| **VII** | Phoenicopterimorphae/Mirandornithes (Flamingos and Grebes) |

Additionally, different analytical methods, and/or the inclusion or exclusion of multiple data types (Reddy et al. 2017) can result in conflicting resolutions, with issues regarding methodology (e.g., inaccurate evolutionary models) may be more difficult to address. In particular, difficult-to-resolve edges may be the result of low signal within very short internodes. Concatenation can overcome low signal by combining the signal from multiple genes but also assumes that all genes share a single evolutionary history, yet population genetic and molecular evolutionary processes can result in highly heterogeneous gene trees and violate this assumption (Kolaczkowski and Thornton 2004; Kubatko and Degnan 2007; Edwards 2009). The most commonly used summary coalescent methods (e.g., ASTRAL III, Zhang et al. 2018; MP-EST, Liu et al. 2010), on the other hand, accommodate gene tree conflicts but assume accurately estimated gene trees for neutrally evolving gene regions that are free of intralocus recombination and only exhibit conflict due to ILS (Springer and Gatesy 2016). However, typical phylogenomic datasets include many genes that violate at least some of these assumptions (although the methods may be robust to these model violation). Also, when analyzing large genomic datasets, many genes will have experienced episodic periods of strong selection, or linkage and hitchhiking can also cause violation of neutral evolution for most genomic regions (e.g., Wang et al. 2021). Moreover, some genes show within-gene conflict, supporting multiple trees (Kimball et al. 2013; Mendes et al. 2019). These observations highlight the importance of examining individual genes in phylogenetic analyses, especially when errors in alignment, assembly, and gene-tree estimation may be common as dataset size increases and larger datasets are more difficult to curate (Gatesy and Springer 2017; Liu et al. 2017).

With ~10,500 living species, birds represent the most species-rich group of tetrapod vertebrates. Despite their importance, several controversial relationships in Neoaves have not been settled (e.g., Reddy et al. 2017). Challenges to resolving these nodes are not unexpected considering that neoavian birds likely experienced a very rapid radiation (e.g., Ericson et al. 2006; Jarvis et al. 2014; Prum et al. 2015; Houde et al. 2020). Although genome-scale datasets have facilitated the recovery of many relationships (e.g., passerines + parrots, flamingos + grebes, sunbittern + tropicbirds, and core landbirds; Hackett et al. 2008; Wang et al. 2012; McCormack et al. 2013; Jarvis et al. 2014; Suh et al. 2015; Braun et al. 2019; Sangster et al. 2022), the Neoaves are still well-known for multiple, extremely recalcitrant nodes. Specifically, the earliest divergences of Neoaves remain one of the most difficult phylogenetic problems in vertebrates, which has even been suggested to represent a hard polytomy (Suh 2016, but see Reddy et al. 2017). Within Neoaves, the positions of Hoatzin *(Ophisthocomus hoazin)* and owls (Strigiformes) remain especially problematic as well as being significant for the analysis of avian development, life history, and morphological traits. The Hoatzin is a bizarre and enigmatic bird species. It converts plant cellulose to simple sugars using microbial foregut fermentation (Domínguez-Bello et al. 1994) and has a modified skeleton (sternum) to accommodate a large crop. Also, Hoatzin chicks have claws on two of their wing digits (Hughes and Baker 1999). The unique features of the Hoatzin have confounded efforts to understand its phylogenetic position for decades, leading to the placement of Hoatzin with diverse taxa such as landfowl (Galliformes), cuckoos (Cuculiformes), or turacos (Musophagiformes) (Hughes and Baker 1999). Owls are nocturnal predators with raptorial features, but their relationships have been obscure. Within Neoaves there are at least two distinct raptorial lineages. The first is the falcons (Falconiformes *sensu stricto)* which have now been shown to be related to parrots and passerines, whereas the second, Accipitriformes, includes the hawks, eagles, and vultures. Presently, it remains unclear whether owls represent a third raptorial lineage or whether they are sister to hawks, eagles, and vultures. This uncertainty impacts our understanding of the evolution of raptorial features.

Recent avian studies have suggested that phylogenetic reconstruction is biased by the source of the data (e.g., exon, intron, or UCE [ultraconserved element], Reddy et al. 2017), which we refer to as data type effects. For example, in Jarvis et al. (2014), concatenated analyses of exons, introns, and UCEs were largely discordant among each other. The resolution of owls and the topology for the earliest split of Neoaves have been shown to vary among different data types (Reddy et al. 2017; Braun et al. 2019; Braun and Kimball 2021). In contrast, the position of the Hoatzin has been even more variable across studies (e.g., Reddy et al. 2017) and its position does not exhibit an obvious correlation with data type. Due to biased base composition and high GC content variation (hereafter GCCV), exons received the most criticism for model violation and impact on phylogenetic reconstruction (Jarvis et al. 2014; Reddy et al. 2017; Braun and Kimball 2021). Using models that incorporate base compositional heterogeneity (e.g., Galtier and Gouy 1998; Morgan et al. 2013) might improve analyses, but dramatic incongruence among gene trees is often observed within data types (e.g., Jarvis et al. 2014) so the available models that incorporate base compositional heterogeneity are unlikely to eliminate incongruence among exonic trees. Nevertheless, finding that exons are generally misleading in the relationships they reconstruct would have far reaching implications for phylogenomic analyses as transcriptomic and targeted sequencing data have become more common (e.g., Prum et al. 2015; Wang et al. 2019).

Herein we examine conflicting signals within two major avian datasets: Jarvis et al. (2014) and Prum et al. (2015). We conduct constrained phylogenetic analyses (edge-based analyses [EBA], see Smith et al. 2020) to examine the relative support for alternative resolutions of three relationships within Neoaves that have proven to be difficult to resolve; specifically, we examine the earliest divergence within Neoaves, the position of Hoatzin, and the position of owls. We explore the efficacy of large datasets to resolving these relationships by carefully curating individual genes and assessing their contribution to the support for alternative hypotheses. In addition to individual gene evaluations, we focus on dissecting the phylogenetic signal associated with different data types (exon, intron, and UCE) and examine other factors, such as GCCV (Foster and Hickey 1999) and gene evolutionary rate (Chojnowski et al. 2008) that might also influence phylogenetic estimation. The inclusion of multiple data types facilitates examination of whether phylogenetic signal from protein-coding regions were different from those of non-coding regions. Through these analyses, we seek to better characterize the ability of large datasets to address difficult phylogenetic questions in birds and other groups.

## 2. Materials and Methods

We have illustrated the data sources and overall analytical framework for this study using the flow chart in Fig. 1.

**Figure 1.**
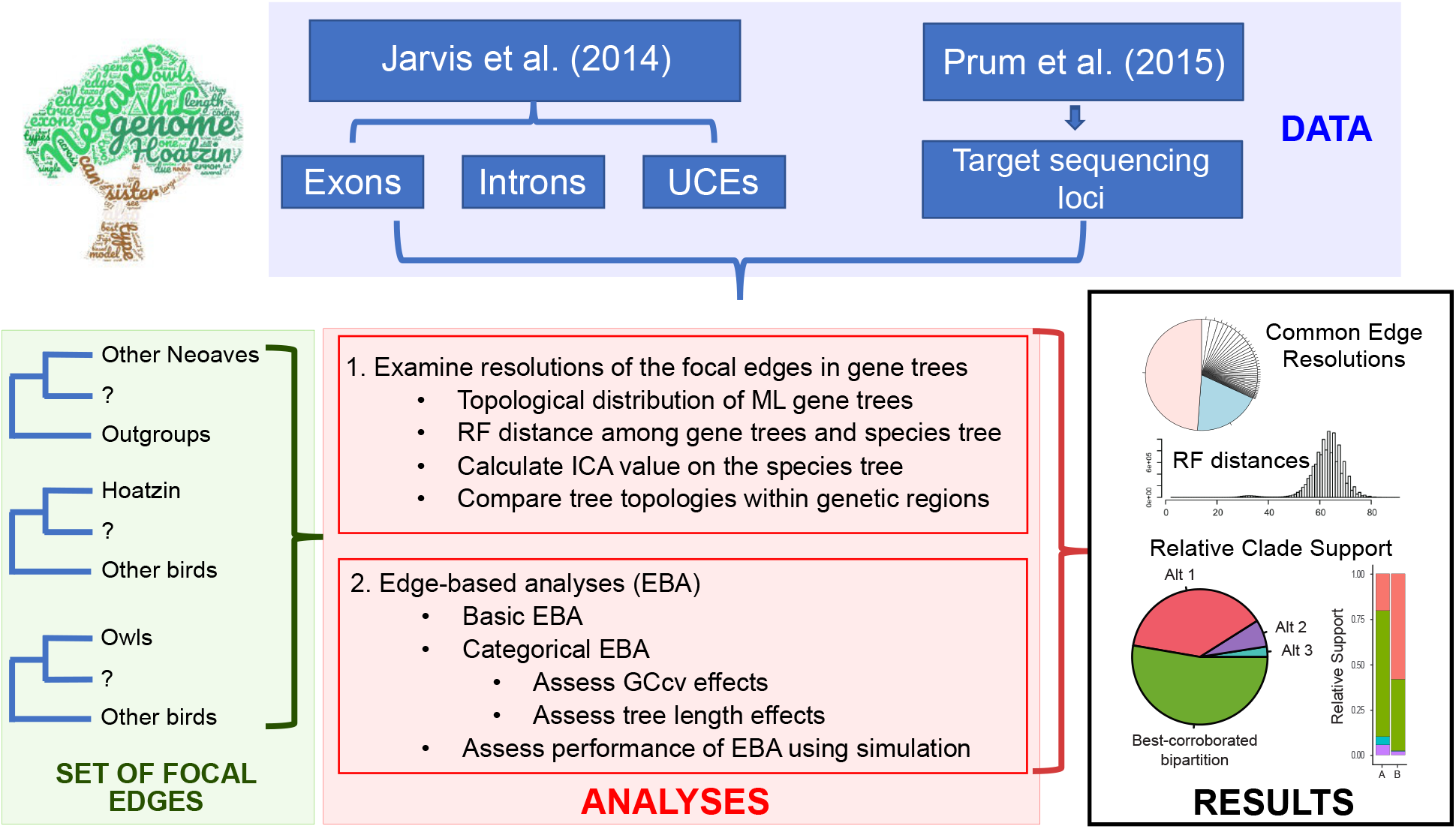
Flow chart illustrating the analytical processes in the current study. Briefly, we used four types of sequence data (blue box, top) obtained from Jarvis et al (2014) and Prum et al. (2015) to examine the support for three selected edges in the avian tree of life (green box, left). Then we conducted two types of analyses using the sequence data (red box, center). First, we examined the resolutions of the three focal edges in gene trees. The results of those analyses (black box, right) were most common resolutions of the edges (upper pie chart) and information about the distribution of distances among gene trees (e.g., histogram of RF distances). Those results, combined with prior information from the literature, allowed us to define the set of plausible resolutions for each focal edge. EBA yields information about the relative support for each of those plausible resolutions; we present the relative support for each resolution either as a pie chart or as stacked column graphs (showing in the lower part of black box). The largest slice in this example pie chart (green) is the best-corroborated relationship, but the chart also shows three alternatives with substantial differences in their relative support. The interpretation of stacked column graphs is similar, but the use of columns makes it easier to compare differences in the relative support for various subsets of the data (in this example, data subsets A and B were compared).

### 2.1 Data acquisition and filtering

We obtained 8251 exons, 2516 introns, and 3769 UCEs from Jarvis et al. (2015) (hereafter Jarvis data). To increase the phylogenetic signal from individual genes and eliminate potential systematic error or noise, we conducted several filtering steps. First, genes having a short alignment (< 500bp) and/or missing taxa (< 42 species after deleting sequences with > 70% gaps in the alignment) were excluded. Second, we reconstructed ML gene trees using IQTREE (v. 1.6.3, Nguyen et al. 2014) with the GTR + Γ model and 1000 Ultra-Fast Bootstrap (UFBS, Minh et al. 2013) replicates. Trees having outgroups that clustered within ingroups (i.e., non-bird species nested within birds, and/or species from Palaeognathae and Galloanserae nested within Neoaves) were treated as systematic error and excluded from additional analyses. Finally, trees with extremely long branch lengths (> 1 expected substitution per site) were also excluded due to potential error in ortholog identification. Scripts are available at https://bitbucket.org/ningwang83/avian_conflict_assess. In addition, the original 259 sequence alignments and gene trees from Prum et al. (2015) were downloaded and filtered following a same procedure (hereafter Prum data).

### 2.2 Examination of gene tree resolution diversity

In general, we focused on three controversial edges on the avian tree of life: the earliest divergence in Neoaves and those encompassing the positions of the Hoatzin and owls (Table 2). We assessed the diversity of resolutions in the gene trees as follows. First, we examined the topological distributions of ML gene trees (Fig. 2) in relation to the three focal edges under different support cutoffs (i.e., ≥ 0%, 50%, 70%, 95% support based on either BS [bootstrap] or UFBS, Hoang et al. 2018). Second, we examined Robinson–Foulds (RF, Robinson and Foulds 1981) distances among gene-trees for each data type, as well as between each gene tree and the concatenated TENT_ExaML species tree from Jarvis et al. (2014) and the concatenated RAxML tree from Prum et al. (2015). RF distances were calculated using the *bp* script (https://github.com/FePhyFoFum/gophy), which allows non-overlapping taxa in the gene trees. Finally, to quantify the relative strength of the phylogenetic signal for a certain edge, we calculated the ICA values (Internode Certainty All; Salichos et al. 2014; see Table 1) on the species trees (same as above) from the filtered gene trees of each data type respectively. These analyses were conducted using RAxML 8.2.11 (Stamatakis 2014) with the GTRCAT model and lossless support to accommodate partial gene trees (Kobert et al. 2016).

**Table 2.** Three focal edges and major testing hypotheses that published before.

| Focal Edges | Hypotheses | Major References |
| --- | --- | --- |
| Earliest Neoaves | 1. **VI+VII** | Jarvis et al. 2014; Reddy et al. 2017; Braun and Kimball 2021 |
|  | 2. **V** | Jarvis et al. 2014, Prum et al. 2015 |
|  | 3. **VII** | Jarvis et al. 2014; Braun and Kimball 2021; Kuhl et al. 2021 |
|  | 4. **III+V+VI+VII** | Hackett et al. 2008 |
|  | 5. Doves | McCormack et al. 2013 |
| Hoatzin's sister | 1. **I+III+V** | Reddy et al. 2017 |
|  | 2. **I** | Prum et al. 2015 |
|  | 3. *S+CR* | Jarvis et al. 2014 |
|  | 4. **V** | Jarvis et al. 2014 |
|  | 5. *S* | McCormack et al, 2013, Jarvis et al. 2014 |
|  | 6. *TB* | Jarvis et al. 2014 |
|  | 7. **II+III+S+CR** | Jarvis et al. 2014 |
| Owls' sister | 1. *M+W* | Jarvis et al. 2014; Prum et al. 2015; Reddy et al. 2017 |
|  | 2. *W* | McCormack et al. 2013; Jarvis et al. 2014 |
|  | 3. *EHV* | Jarvis et al. 2014 |
|  | 4. *M* | Hackett et al. 2008; Gilbert et al. 2018 |

NOTE.— the Greek numbers (bolded) correspond to the “magnificent seven” of Reddy et al. (2017), with other italic acronyms that can be referred to Table 1.

**Figure 2.**
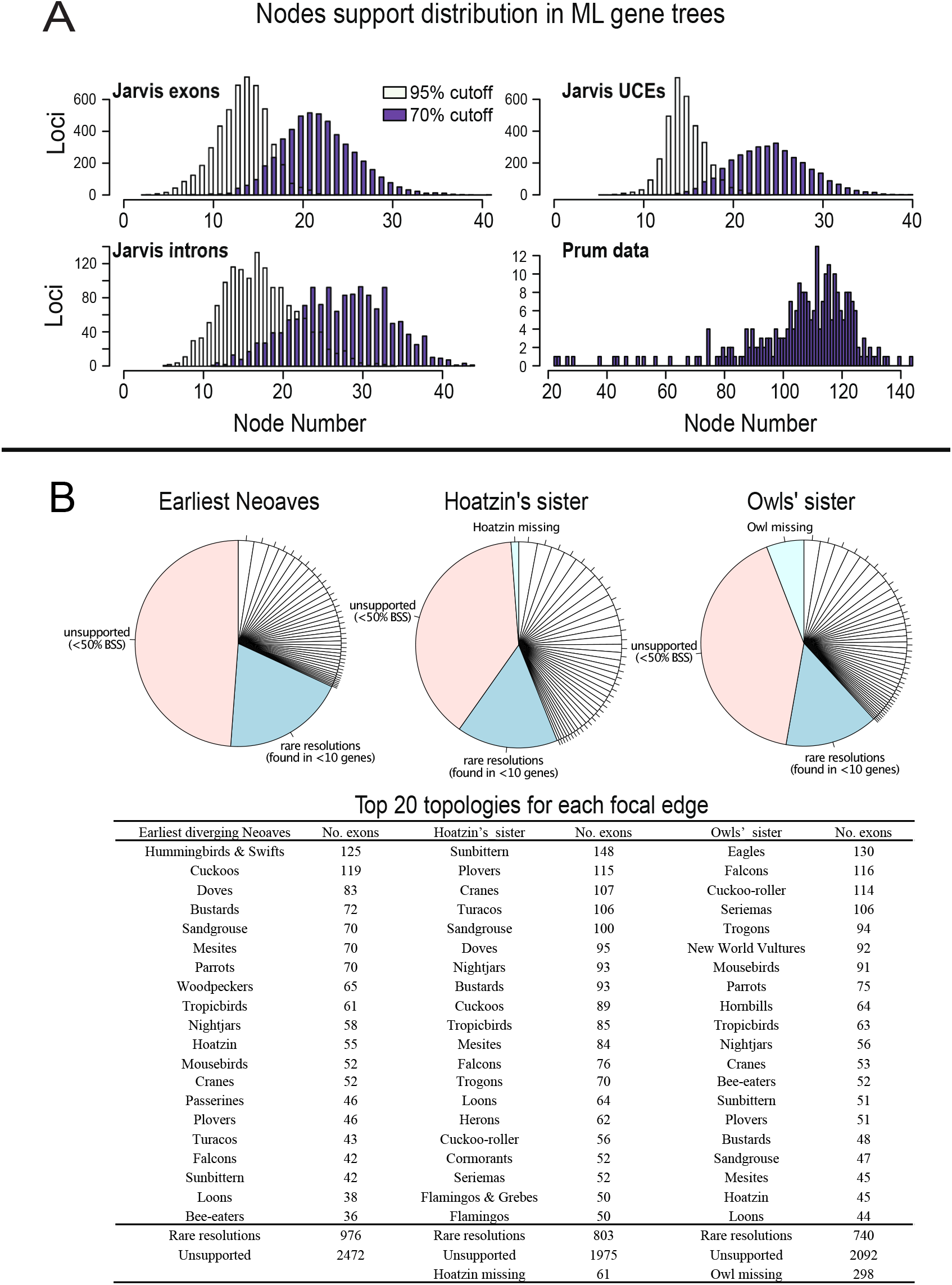
A: Node support distribution among ML gene trees. B: Topology distribution of the ML gene trees for the three focal edges using Jarvis et al. (2014) exon data. Analyses for Introns and UCEs can be found in Figure S3. unsupported: relationships shown have less than 50% UFBS (ultrafast bootstrap support, we use lower support cutoff to show potential topology distributions). The Top 20 supported topologies are shown. rare resolutions: relationships shown in less than 10 gene trees. Hoatzin missing or Owl missing: Hoatzin or Owl is missing in the gene trees.

### 2.3 Edge-based analyses — Assessment for constraint topologies for three focal edges

We selected several alternative topologies from previous large-scale studies (Table 2) for each of the three focal edges. For the Jarvis data, gene trees were reconstructed in IQTREE using one constrained edge at a time and optimizing the remainder of the tree. This analysis was conducted for each locus and every focal edge (Fig. 3). Since the ML gene trees typically exhibited topologies different from published dominant relationships (see discussion bellow), genes were excluded if their ML gene tree was incompatible with any testing constraint under a 95% UFBS cutoff (i.e., any locus with an ML gene tree that has a focal node conflict with all test hypotheses would be kept only if the conflicting node was weakly supported [< 95% UFBS]). To identify the best hypothesis, we conducted the following analysis:

1. We compared *lnL* (log likelihood) scores among conflicting alternative topologies for a single edge for each gene and identified its supporting (i.e., the best) topology.
2. We calculated the difference in ln*L* (Δln*L*) between the best and the second-best resolutions for each gene. At this stage, we also identified potential outlier genes (cases where Δln*L* fell out of the smooth distribution among other genes) by eye.
3. We summed Δln*L* for each constraint from supporting or significantly supporting (Δln*L* > 2) genes (referred to as ∑Δln*L* or ∑sigΔln*L*).
4. We identified the overall best topology either supported or significantly supported (Δln*L* > 2) using two criteria: the largest number of genes (∑Ng: simply the number of genes supporting a specific topology) and the highest ∑Δln*L* (or *∑*sigΔln*L*) scores (Fig. 3).

**Figure 3.**
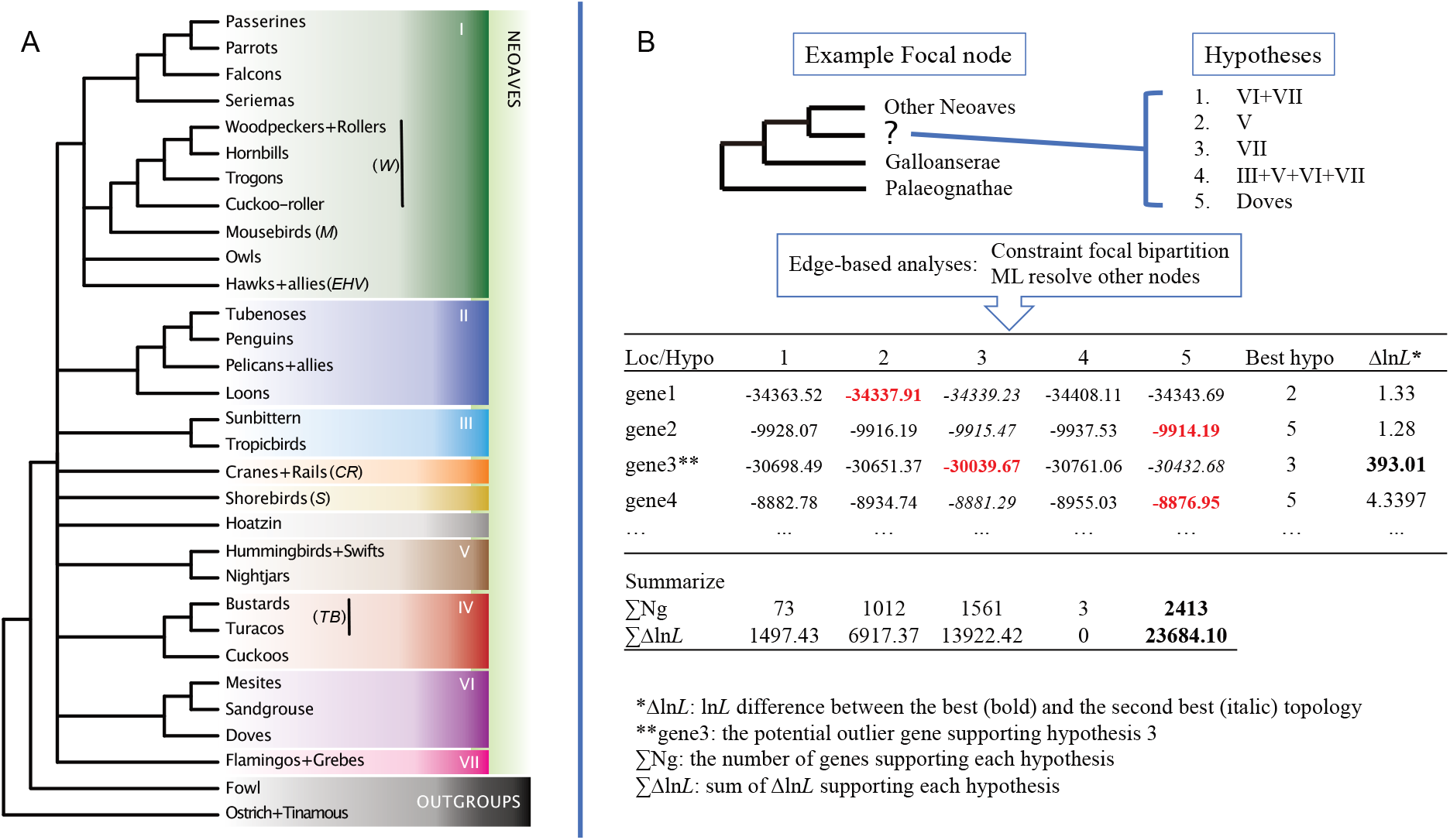
A: Keys (**I-VII**) for each major clade in the hypotheses being tested. B: flow chart on how to conduct edge-based analyses by using the earliest diverging Neoaves as an example.

Genes for which the *lnL* could not be calculated given specific constraint topologies (this occurs when relevant taxa are missing) were excluded from the calculation.

We manually examined the outlier genes for misalignment and collected information about the functions of them. We did this because positive selection can give rise to convergent evolution, at least in principle, and genes with a specific function might group corresponding species together and mislead phylogenetic inferences. For example, the hearing gene *Prestin* grouped echolocating taxa (microbats and toothed whales), presumably reflecting convergence (Li et al. 2010). Using BLAST, we determined outlier exons whether their function might be related to the outlier topology. However, we did not conduct lineage-specific selection analyses for outlier exons as their alignments contained several premature stop codons.

To control for the impact of taxon sampling on topological preference, we also conducted EBA using genes from the large sampling (i.e., 198 bird species) of Prum et al. (2015), following similar processes and criteria indicated above. To limit CPU time, the RAxML 8.2.11 (Stamatakis 2014) with GTRCAT model was used. The ML gene-trees including BS support built with RAxML were directly retrieved from Prum et al. (2015). We also reduced the number of constraint topologies to include only three hypotheses for each focal edge (see Results and Discussion 3.3). Results were organized after filtering out loci with gene-tree incompatibilities with any testing constraints under 70% BS cutoff (Hillis and Bull 1993). In addition, to examine the sampling effect on the hypotheses examined here, we focused on the earliest diverging Neoaves and assessed five additional hypotheses by referring one single lineage (i.e., caprimulgiforms [nightjars], Hoatzin, charadriiforms [represented by plovers], gruiforms [cranes], or cuculiforms [cuckoos], Fig. 4) as sister to other Neoaves and conducted EBA with Jarvis data using the same criteria mentioned above.

**Figure 4.**
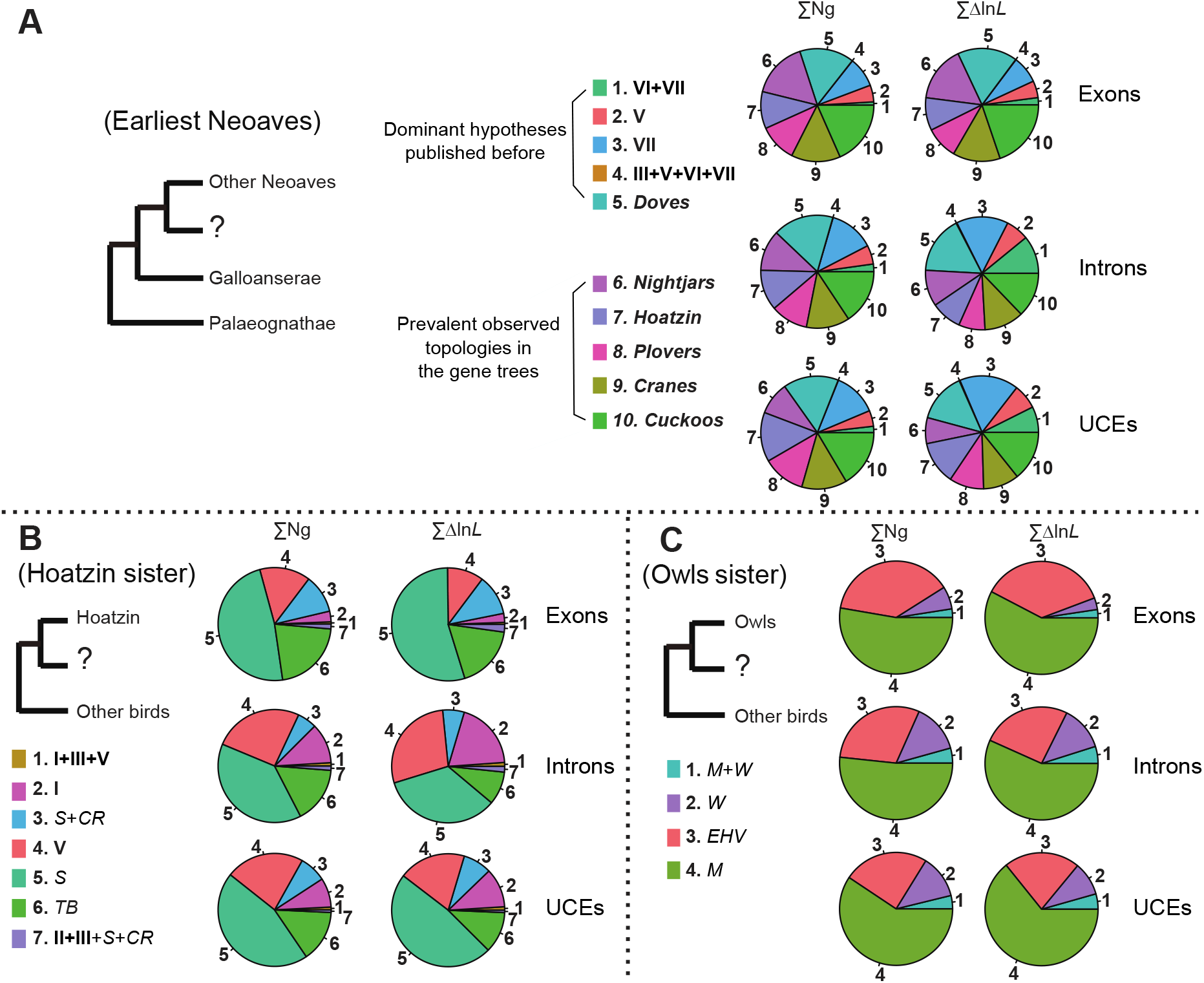
The three focal edges and the testing hypotheses in the edges-based analyses using exons (upper), introns (middle) and UCEs (lower). A: The earliest diverging clade in Neoaves, including the five dominant hypotheses published before (1–5), and the five additional hypotheses prevalent observed in the gene trees for bias assessment (6–10). B: the sister of Hoatzin, including seven previously identified hypotheses. C: the sister group of Owls, including four commonly found hypotheses. ∑Δln*L*: the sum of log likelihood score differences; ∑Ng: the number of genes supporting a specific hypothesis. Hypotheses keys can be found in Figure 3.

In order to assess influence of GCC_V_ and tree length (i.e., gene evolutionary rate) on tree reconstruction using different data types, we sorted the filtered genes from Jarvis et al. (2014) based on their GCC_V_ and tree length. We characterized GCC_V_ using a method similar to Reddy et al. (2017). We excluded the crocodilian outgroups and calculated the GC content for sites that are variable within birds. Then we calculated three different ΔGC metrics (ΔGC_50_, ΔGC_90_, and ΔGC_100_), which represent the GC content difference between a high-GC and a low-GC taxon. The three different metrics differ in the high- and low-GC taxa that we selected; for ΔGC_50_ it was the upper and lower quartile, for ΔGC_90_ it was the 95^th^ and 5^th^ percentile, and for ΔGC_100_ it was the maximum and minimum. We ordered genes based on ΔGC_90_ to balance the GC difference and similarity among data types (i.e., maximize differences while eliminating outlier signals, Fig. 5A) and examined the gene trees in the upper (i.e., high GC_CV_) and lower (i.e., low GC_CV_) quartiles of the variation for their supporting pattern to each testing edge (Fig. 5B). We conducted similar analyses by sorting genes based on their total tree length and compared support for focal edges between gene sets from the upper and lower quartile of the tree lengths (Fig. 5C and D).

**Figure 5.**
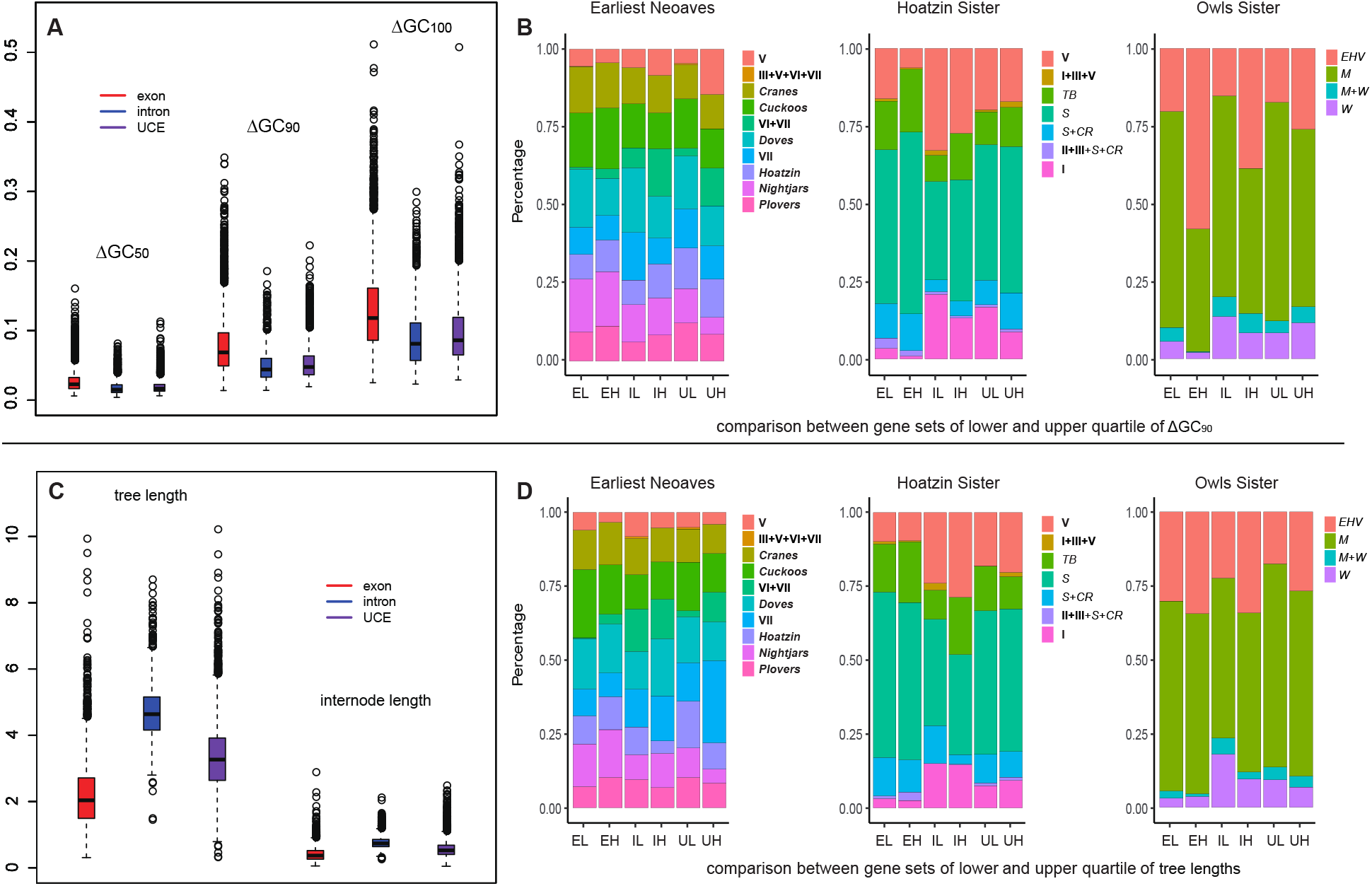
A: The GC content variation among exons, introns, or UCEs, including ΔGC_50_, ΔGC_90_, and ΔGC_100_ (see Table 1 for explanation). B: Comparison of hypotheses supporting pattern between gene sets from the lower and upper quartile of ΔGC_90_ among data types. C: Tree length (left) and internode length (right) variation among different types of data. D: Comparison of hypotheses supporting pattern between gene sets from the lower and upper quartile of tree length variation. For x axis, EL/EH: exons with lower or higher ΔGC_90_/tree length; IL/IH: introns with lower or higher ΔGC_90_/tree length; UL/UH: UCEs with lower or higher ΔGC_90_/tree length.

### 2.4 Comparison of tree topologies within genetic regions

Within-genetic region comparisons were conducted using exons and UCEs from Jarvis et al. (2014), which have high alignment quality without long indels, in addition to the Prum dataset. We split the genes into 1000bp chunks, omitting the short leftovers. We conducted ML analyses in IQTREE based on GTR + Γ model with uncertainty measured using both SH-aLRT and UFBS, each with 1000 replicates. Edges with < 80% SH-aLRT or < 95% UFBS support were collapsed. We note that this requirement is extremely conservative but limits false positives. We also reestimated gene trees using these same criteria for consistency among these tests. We then compared trees from gene region fragments to the ML gene trees based on the entire gene region and noted conflicts.

### 2.5 Coalescent simulation – assessing the performance of EBA for deep coalescence

We conducted simulations to assess the performance of EBA in recovering the potential true topology at edges under differing level of ILS. The Jarvis et al. (2014) MP-EST* TENT tree, which was built with a binned MP-EST* method (Mirarab et al. 2014), was used as the *model species tree*. We rescaled the terminal branch length of the model species tree to make it ultrametric and generated 500 simulated gene trees using the neutral coalescent model with the *treesim.contained coalescent tree* module in Dendropy 4.2.0 (Sukumaran and Holder 2010). We rescaled the internal branches of these gene trees with 0.01, which is estimated by comparing the coalescent and substitution branch lengths (BLs) in the empirical data of Jarvis et al. (2014). Duchêne et al. (2018) found that the tip branch-lengths (tipBLs) are best to be proportional among species across gene trees. Thus, we first obtained the distribution of BLs for each species from the empirical ML gene trees of Jarvis et al. (2014). Then, we generated tipBLs for each simulated gene by selecting the same percentile tipBLs value from the empirical distribution of each species. This rescaling method ensures a proportional tipBLs across all gene trees. To simulate different types of data, we estimated the branch length distributions, and the modeling parameters (see below) from exons, introns and UCEs, respectively. Here for each datatype, we built a coalescent species tree with ASTRAL III (Zhang et al. 2018) using these rescaled simulated gene trees. Moreover, the average RF distance (ARFD) between the model species tree and the simulated gene trees was used to reflect the level of overall ILS.

We generated one sequence alignment with length of 800-3000bp for each simulated gene tree using Seq-Gen (Rambaut and Grass 1997, implemented in Dendropy). All alignments were generated using the GTR + Γ model with model parameters (base composition, α parameter for Γ distribution, and GTR substitution rates) selected randomly from the density distribution of the Jarvis data of exons, introns, and UCEs, respectively. Simulations were repeated once for each data type to validate results, so there were six sets of 500 simulated alignments in total. Then we obtained *ML estimates of simulated gene trees* for these simulated alignments in IQTREE using the GTR + Γ model. In addition, 184 bipartition constraints were generated from 27 species-tree estimates constructed by Jarivs et al. (2014), which provide a reasonable representation of the extent variation of species tree hypotheses as to bird relationships, so that all the bipartitions in the model species tree were included. The EBA were conducted using the simulated alignments following the method described above on the three difficult focal edges that exhibit higher level of ILS as well as some commonly accepted edges that exhibit lower levels of ILS, including **I**, **II**, **III**, and **V** (Table 1) from the “magnificent seven” superordinal groups (Reddy et al. 2017). We also conducted ASTRAL analyses using the ML estimates of simulated gene trees. We calculated the mean gene-tree estimation error (GTEE) by using Dendropy to calculate the unweighted RF distance between the simulated gene trees and the ML estimates of the simulated gene trees, which we averaged across all genes. To compare the performance of concatenation analyses, we concatenated the simulated alignments of each data type and conducted ML analyses in RAxML 8.2.11 using GTR + Γ model, partitioned by locus, and we calculated node support using 200 bootstrap replicates.

## 3. Results and Discussion

### 3.1 Gene informativeness

After careful data curation, we obtained 5062 exons, 1409 introns and 3370 UCEs (Table S1) from Jarvis et al. (2015). All data from Prum et al. (2015) passed the filtering processes and were therefore retained.

We examined the extent to which each genomic partition had information to resolve nodes within the tree. We found that each gene in the Jarvis dataset supported, on average, 24 (≥ 70% UFBS) or 15 (≥ 95% UFBS) of the 47 internal nodes, respectively, whereas the Prum data supported (≥ 70% BS) ~110 out of 197 internal nodes on each gene tree, with a maximum of 144 and minimum of 22 (Fig. 2A). The low support on many nodes is likely caused by the relatively low phylogenetic signal for individual genes that are subject to high GTEE (Molloy and Warnow 2018), especially across such broad relationships. Although introns generally supported more relationships, the number of supporting nodes appeared to correspond to alignment length (Fig. S1). For the three edges of interest, only 23%— 33% exons, 38%—51% introns, and 34%-43% UCEs in the Jarvis data obtained ≥ 70% UFBS on ML gene trees after excluding genes with missing owls or Hoatzin. In the Prum data, similarly, only four loci obtained supports (>70% BS) on their ML gene trees for the resolution of Hoatzin, two genes showed supports for the earliest diverging clade of Neoaves, whereas no gene supported the position of owls. 113 genes even showed non-monophyly of owls (i.e., the two families of owl had distinct positions in the tree) in Prum et al. (2015). These results suggest that most loci have limited information for resolving the contentious edges and therefore a high potential for GTEE from these genome data, regardless of the sources of conflict.

### 3.2 Diversity of resolutions

Many gene regions across datasets and datatypes did not demonstrate a strong preference for any specific resolution regarding the three focal relationships (Fig. S2). However, to ascertain the nature of the support for alternative resolutions when support was indicated, we firstly examined how many different resolutions were supported by different genes. We observed, for example in the Jarvis exons, there were more than 1500 alternative resolutions for each focal relationship. Most of these were unsupported (~ five resolutions obtained ≥ 95% UFBS and ~40 resolutions obtained ≥ 50% UFBS) or were present in fewer than 10 gene trees (i.e., rare resolutions, Fig. 2B). Even the best-supported topology for each data type was still present in less than 5% of gene trees. The situation was similar for introns and UCEs (Fig. S3A). The Prum data had 259 loci, and we identified 110 or more topologies for each focal edge, but only 3 – 16 topologies had > 50% BS, within which more than 80% were found in a single gene (Fig. S3B).

To properly discuss the uncertainty of the focal relationships and the large number of plausible topological alternatives, we have adopted a compact notation for clades. For example, the root of Neoaves in the Jarvis et al. (2014) TENT_ExaML is between **VI** + **VII** (Table 1) and all other Neoaves. Hereafter, we refer to this topology as **VI**+**VII** sister, and both the **VI**+**VII** clade and the remaining Neoaves (i.e., the edge that supports the **VI**+**VII** sister relationship) have 100% BS in the TENT_ExaML and the binned MP-EST* trees in Jarvis et al. (2014). Houde et al. (2019) conducted additional analyses of the Jarvis non-coding (intron and UCE) data, using both nucleotide sequences and indels, and they also reported the same basal topology (albeit with local posterior probabilities <1.0). However, the **VI**+**VII** sister obtained −0.051 (intron), −0.272 (UCE), and −0.317 (exon) ICA values in our analyses, indicating substantial conflicting signals among gene trees, and a better supported alternative conflicting relationship as shown in the ML topology distribution (Fig. S3). The very different ICA values are also indicative of a data type effect, with introns showing a very different signal than UCEs or exons. Similar situations were also found for the other two focal edges (i.e., for the Hoatzin and owls) and in both datasets (not shown).

The diversity of resolutions can also be observed by the RF distances among gene trees. Trees based on Jarvis exons and UCEs exhibited more among-gene tree differences (~65, 69% of the edges, Fig. S4) than do trees based on introns (~58, 62% of the edges), implying the potential of high GTEE in these gene trees (Simmons et al. 2016). Similar patterns were also found when comparing gene trees to the species-tree topologies. The RF distances among gene trees for the Prum data were usually larger than 150 (38% of the edges) and as high as 300 (76% of the edges, Fig. S4), suggesting high topological diversity that may be a reflection of dense taxon sampling in the data.

### 3.3 Edge-based analyses (EBA)

Examining conflicting signals in large-scale datasets is perhaps the foremost challenge in systematics as we have entered the phylogenomic era. One common analytical framework is concatenation, which is known to be a useful methodology in many contexts (Bryant and Hahn 2020) despite its failure to model the multispecies coalescent (Edwards et al. 2016). However, the utility of concatenation for the examination of conflict is limited because they generally yield a single tree. Low support values in concatenated analyses can either provide evidence of conflict or it could reflect the absence of any signal in the data. Fundamentally, there are only two ways to examine conflict using the concatenated approach. First, one can subdivide data based on some criterion, such as coding vs. non-coding data (e.g., Jarvis et al. 2014, Reddy et al. 2017; Braun and Kimball 2021) or protein structure (e.g., Pandey and Braun 2020; Gordon et al. 2021), and conduct concatenated tree searches on the data subsets. Second, one can examine differences in the tree selection criterion (typically the likelihood) for various topologies at the level of the genes (Shen et al. 2017) or the individual site level (Kimball et al. 2013). Both approaches have limitations: the former only allows assessment of data type effects whereas the latter can only yield information about a small number of topologies due to computational time limit.

An alternative way to examine conflicts involves comparisons of gene trees that can have any topology. However, analyses of individual genes are subject to high GTEE, so many bipartitions in estimated gene trees may be spurious. Moreover, there can be conflicts within gene regions due to intralocus recombination and other factors (see section 3.5). Ultimately, it is difficult to determine whether the extensive conflict among estimated gene trees reflects a mixture of meaningful biological signals or simply random resolutions due to GTEE. Our EBA of genes lie between concatenation and simple gene-tree analysis. This approach limits the consideration of gene-tree topologies to those containing a specific bipartition. In this study, we have limited the testing bipartitions to those commonly identified previously (see Table 2) or those present in large numbers of gene trees. Comparison of the results given the two complementary criteria, ∑Δln*L* and ∑Ng, can also provide evidence for outlier genes; when ∑Δln*L* for a specific edge is much larger than ∑Ng, it might imply that a relatively small number of genes are contributing a large amount of the ln*L* differences among alternative resolutions. Overall, this edge-based approach should limit GTEE while still retaining the advantages of gene tree-based analyses and provide information about conflicting signals (see also discussion in Mount and Brown 2022).

We examined gene-wide support for the dominant alternative resolutions of the three focal edges that have been previously published (Table 2). After excluding loci with strong incompatible node to our test hypotheses, we obtained 4194 — 5019 exons, 1134 — 1274 introns, and 3165 – 3295 UCEs for each focal relationship (Table S1) from the filtered Jarvis data.

Regarding the earliest diverging Neoaves (Fig. 4A and Table 2), all data types had the highest support for placing doves sister to all other Neoaves (Fig. S2) when five published hypotheses were tested. When we expanded the set of hypotheses to include the most prevalent topologies in the gene trees (Fig. 4A, hypotheses 6-10), support was more evenly distributed across them (Fig. 4A). For the other two edges, Hoatzin were sister to shorebirds *(S),* and owls were sister to mousebirds *(M,* Fig. 4B and C). These findings obtained support, although few, in previous studies, including owls + *M* in a reanalysis of shallow-filtered Jarvis UCE data (Gilbert et al. 2018), doves sister to all other Neoaves in the UCE indel tree (Houde et al. 2019), and Hoatzin + *S* in the complete UCE data matrix (McCormack et al. 2013). Moreover, the relationships published in the Jarvis et al. (2014) TENT tree (i.e., **VI+VII** as earliest Neoaves, *S+CR* sister to Hoatzin, and *M + W* sister to owls) were only weakly supported (Fig. 4).

To examine taxon sampling effect in EBA, we obtained up to 258 loci (Table S2) from Prum et al. (2015) after excluding strong incompatible loci. We tested three constraints for each focal edge, with one constraint directly retrieved from Prum et al. (2015) tree and two from the top two published hypotheses supported by the Jarvis data (i.e., **V/VII**/doves as earliest Neoavian branch, **I**/*S*/**V** sister to Hoatzin, and *[M+W]/M/EHV* sister to owls). The best relationships supported by the Prum data were the same as found in the Jarvis data (e.g., Hoatzin + *S*; owls + *M*, Table S2), indicating a limited taxon-sampling effect on EBA, which increase congruence between analyses of the Prum and Jarvis data. Moreover, analyses focused on the earliest diverging Neoaves using the Prum data revealed a major impact of genes with a large Δln *L*. For example, we noted that gene L111 encoding an FH2 domain containing 1 protein (FHDC1, whose homolog may play a role in both cell shape and microtubule polymerization in humans), disproportionately contributed 171 Δln*L* to support **VII** as the earliest divergence of Neoaves. We did not detect alignment error or lineage specific positive selection for this gene. Moreover, eliminating FHDC1 would not shift the best supported hypothesis given the ∑Δln*L* criterion (Table S2). Regardless of the details, the best-supported hypothesis in these EBA of the Prum data were never the relationships present in the Prum et al. (2015) tree; they were hypotheses found in EBA of the Jarvis data.

### 3.4 Complex data type effects revealed by categorical edge-based analyses

The relatively even distribution of support for various hypotheses regarding the earliest diverging Neoaves when we examined 10 alternatives (Fig. 4A) suggests a very high degree of conflict among gene trees. However, this result also suggests a hypothesis-sampling-effect (i.e., the more hypotheses tested, the more widely distributed the support). It further provides evidence for a data type effect on best hypotheses: exons supported cuckoos sister, introns continued to support doves sister, and UCEs supported either clade **VII** (largest ∑Δln*L*) or cuckoos (largest ∑Ng). It is noteworthy that Jarvis et al. (2014) placed **VII** sister to all other Neoaves in their concatenated analysis of UCEs, and the *∑ΔlnL* criterion is similar to the criterion used for tree selection in ML analyses. However, the observation that UCEs placed **VII** sister in fifth place based on the alternative criterion – ∑Ng, suggests that the support for **VII** sister reflects a limited number of UCE loci with large Δln*L* values (Fig. 4A, Table S3). However, the functions of many UCEs are unclear and it is difficult to examine selection on non-coding loci, so we did not attempt to determine whether there might be a functional basis for the outlier UCEs. There were other differences among data types in the rank order of suboptimal hypotheses. For example, when the ∑Δln*L* criterion is used, **VI + VII** as the earliest Neoavian branch received much higher support in analyses of non-coding data, especially introns, than it did in analyses of coding data (10.9% for introns, 7.4% for UCEs, and 2% for exons). These results agree with the data type effects reported by Reddy et al. (2017) and by Braun and Kimball (2021), although the differences among data types also appear to be more complex than the results suggested before.

Reddy et al. (2017) pointed out that data type had a major impact on the topology for the deepest divergence in Neoaves, with analyses of coding data supporting a **V** sister topology and analyses of non-coding data supporting either **VI+VII** sister (introns) or **VII** sister (UCEs). Several studies (Jarvis et al. 2014; Reddy et al. 2017; Braun and Kimball 2021) pointed out that avian coding data typically exhibit greater GCC_V_ than do non-coding data. Previous studies (e.g., Jermiin et al. 2004; Duchêne et al. 2017) have demonstrated that high GCCV, when fit to a homogeneous model, can results in biased phylogenetic estimation. Applying models with more heterogeneous assumptions (e.g., Galtier and Gouy 1998; Holland et al. 2013) has the potential to improve estimates of phylogeny, although the available programs that implement these models are challenging to apply to phylogenomic datasets (see additions discussion in Reddy et al. 2017). Since our EBA provide a sensitive tool to examine conflicting signals in phylogenomic datasets we assessed the impact of GCC_V_ in the Jarvis et al. (2014) data on the conflicts among data types that we observed.

Generally, ΔGC_50_ was similar among data types but ΔGC_90_ and ΔGC_100_ variation was larger for exons than for introns and UCEs, consistent with findings in Reddy et al. (2017) (Fig. 5A). After sorting genes based on ΔGC_90_, about 1266 exons (E), 352 introns (I) and 842 UCEs (U) were retained in the upper (H) or lower (L) quartile category. If GC_CV_ has an impact on inference we would expect different patterns of support between GC_CV_ categories within the same data type. Indeed, owls + *EHV* showed much higher support (both ∑Ng and ∑Δln*L*) in all high GC_CV_ makers (Figs. 5B and S5), and such signal is strongest in EH and IH (i.e., the high GC_CV_ exons and introns). It seems the characteristics of high GC_CV_ genes tend to correlate with owls + *EHV.* Braun et al. (2019) suggested that the position of owls in concatenated analyses was subject to a data type effect, with analyses of coding data tending to yield the owls + *EHV* topology. We found that support for owls + *EHV* was similar for the EL, IL, and UL datasets (low GCCV exons, introns, and UCEs, respectively) but much higher in the EH, IH, and UH (the high GCCV datasets). However, the signal for owls + *EHV* was still stronger in the high GCCV exons than in the high GCCV introns and UCEs. Thus, our results imply that an interaction between higher GC_CV_ and data type – not coding vs non-coding data type *per se* – is correlated with the owls + *EHV* signal.

As previously noted, the position of Hoatzin has been highly variable across studies and data type effects have not been noticed before (reviewed by Braun et al. 2019). The dominant signal in our categorical EBA was Hoatzin + *S*, which is also present in the Jarvis exon c123 tree and the McCormack et al. (2013) UCE tree. In the categorical EBA, there was higher support for Hoatzin + *S* in high GC_CV_ markers (compare EH vs EL, IH vs IL, and UH vs UL; Fig. 5B), but the support differences are quite trivial. In fact, support for alternative relationships of the Hoatzin show more variations among data types (Figs. 5B and S5) regardless of GC_CV_ and tree length. The most notable difference was a weaker support for Hoatzin + *S* and a stronger support for Hoatzin + **V** in the introns (Fig. 5B and D). Provocatively, Hoatzin + **V** is present in the Jarvis intron tree (albeit with only 70% BS) but it is not in the Jarvis UCE tree (which included Hoatzin + *S* + *CR*, like the Jarvis TENT). These results are not consistent with a simple coding vs noncoding effect for two reasons. First, the signal for Hoatzin + *S* is very similar in exons and UCEs (Fig. 5B) but the Hoatzin + **V** was elevated in both noncoding data types. Second, Hoatzin + **I**, which is found by Prum et al. (2015) using mostly coding data, received higher support in introns and UCEs (Figs. 5B and S5) than it did in the Jarvis exons (Fig. 5). Thus, in agreement with many previous studies, we find that the position of the Hoatzin is very difficult to resolve. However, our EBA revealed a previously unappreciated and relatively complex data type effect and further showed that Hoatzin + *S* was the strongest signal in all three data types.

The most prominent data type effect hypothesized by Reddy et al. (2017) was the **VI + VII** sister to all other Neoaves (their non-coding topology) and **V** sister to all other Neoaves (their coding hypothesis). Using our categorical EBA, **V** as sister to all other Neoaves did not appear to have better support in exons than in introns or UCEs (Figs. 5B and S5). The data subset with the highest support for **V** as the earliest neoavian branch was the high GC_CV_ UCEs (UH in Fig. 5B). This support for **V** sister appears to result from one outlier gene (chr1_28895_s) that contributes almost 40% of the ∑Δln*L* (Table S3) for this relationship. Unsurprisingly, the total number of genes (∑Ng) that support **V** did not vary much among GC_CV_ categories or data types (Fig. S5). As described above, **VI+VII** did appear to be better supported in introns and UCEs, especially in the data subsets with high GC_CV_ and high tree lengths (i.e., IH and UH, Fig. 5B and D). However, the number of genes (∑Ng) supporting **VI+VII** did not vary much among categories and data types (Fig. S5). Thus, the higher support for **VI+VII** sister appears to reflect a relatively small number of genes that make high Δln*L* contributions; given the much higher proportion of signal for **VI+VII** sister in the IH and UH subsets (Fig. 5B), these genes with a high Δln*L* contribution must be especially common in the high GC_CV_ data subsets.

Evolutionary rate is another major factor in phylogenetic resolution (Chojnowski et al. 2008). Tree length (sum of branch lengths) variation suggests different overall rates of evolution and different data types exhibited different patterns, with introns having larger tree length/internode lengths and less variation than UCEs and exons (Fig. 5C). Tree length in these datasets is primarily the result of tip lengths (Fig. 5C) as a consequence of sampling at high taxonomic levels. In general, tree lengths did not have an obvious impact on support for different hypotheses for each focal edge. However, we noticed two interesting patterns with respect to the earliest diverging Neoaves (Fig. 5D). First, **VI+VII** as sister to all other Neoaves usually received higher support in categories with high GCCV and high tree lengths, but we noted that **VI+VII** also received high support (i.e., the best) in the intron subset of low tree lengths (IL, Fig. 5D). Within this category, we identified one gene (e.g., 11117.intron, Table S3) contributing ~21% of the ∑Δln*L* supporting this relationship. We did not detect any alignment issue in this gene and its tree length is around the upper edge of this quartile set. Thus, its outlier signal may be driven by some biological characteristics that may not be directly related to evolutionary rate. Second, **VII** as sister to all other Neoaves received the highest support from the UCE subset of high tree length (UH, Fig. 5D), consistent with the Jarvis et al. (2014) UCE tree. Again, an outlier gene (chr8_1881_s, that contributed 1/3 ∑Δln*L*, Table S3) was identified, and its support for **VII** as sister may be driven by certain characteristics that may or may not be directly related to its high tree length.

As found previously, genes that have disproportionally high support for specific relationships (i.e., outlier genes) can bias phylogenetic reconstruction (Kimball et al. 2013; Shen et al. 2017). We noted several potential outliers within these datasets (Table S3). As indicated above, some outliers can significantly support certain hypotheses. However, removal of most single outlier genes did not change the results significantly nor could we detect any alignment errors with these genes. The diversity of alternative hypotheses noted earlier made it difficult to detect the underlying reasons for biased support from outliers. One biological process that might cause biased phylogenetic resolution is selection. However, we could not test selection as most exons contained premature stop codons (e.g., single mutation likely caused by sequencing or assembly error). Both introns and UCEs also contain outliers. Although it was impossible to link gene specific traits to the topology they support without annotation, our categorical EBA indeed implied that the earliest diverging taxa in Neoaves might be influenced by outlier signal. In contrast to the case for the earliest diverging Neoaves, we did not detect as strong of an impact from outlier genes on the other two focal edges (the positions of Hoatzin and owls).

### 3.5 Within gene region conflict

To determine whether gene tree conflict was the result of conflicting signal within genomic regions, we further examined phylogenetic signal across regions within each gene. The average gene-region length of the Jarvis data was 1998 bp (501 – 15819 bp) for exons, 2509 bp (1787 – 3527 bp) for UCEs, and s 9309 bp (506 – 66,762 bp) for introns. For Prum data, the average gene region length was 1524 bp (361 – 2316 bp). We focused on exon and UCE alignments as they were smaller and therefore less likely than introns to suffer from mixed signal. We conducted analyses on each gene region to determine whether 1000 bp segments conflicted with the ML tree calculated for the entire region. We found extensive conflicting signal after collapsing edges with < 95% UFBS and < 80% SH-aLRT support. Although this is an extremely conservative measure, and likely to have a high false-negative rate (Guindon et al. 2010), it was chosen to prevent false positives. For loci with at least 2 × 1000 bp segments, we found that 10% of Jarvis et al. (2014) exons (195 of 1985), 6% (207 of 3367) of UCEs, and 50% (8 of 15) of gene regions in Prum et al. (2015) had conflicts. It is noteworthy that exons in Jarvis et al. (2014) usually concatenated multiple exon regions of one gene and that some of those exons are separated by long introns. Thus, those exons might exhibit different evolutionary histories due to recombination, selection, and/or other reasons. Despite the complexity of the Jarvis exons, the number of conflicts we observed in this analysis was only slightly higher than it was for UCEs. Although intragenic conflicts within the Prum data were much higher, they could reflect sampling error due to the small number (i.e., only 15) of long gene regions in the data; the larger number of taxa in the Prum dataset is also likely to have had an impact.

To determine whether within gene conflict corresponded to obvious data “dirtiness”, we examined alignments for gene regions that exhibited conflict. All large datasets suffer from some amount of noisy data (Springer and Gatesy 2018; Braun et al. 2019) and we did find some cases where intragenic conflicts might reflect misalignment. For example, the 1424.exon gene region from Jarvis et al. (2014) had an obviously “messy” alignment (Fig. S6); analyses of problematic segments placed Rifleman sister to pelicans, mesites sister to cranes, grebes sister to sandgrouse, and fulmars sister to seriemas. However, many other loci that exhibited substantial intragenic conflicts had alignments that appear largely unambiguous (e.g., locus 9838.exons, Fig. S6), which suggests potential biological processes, like those highlighted by Scornavacca and Galtier (2017) and Mendes et al. (2019), underlie the observed intragenic conflicts.

### 3.6 Simulations

Most of the gene-tree topologies resulted in a single species/lineage as sister to the remaining lineages (Figs. 2B and S3). To determine if this may be expected given that short internodes should have a higher prevalence of ILS, we conducted coalescent simulations using the MP-EST* TENT tree from Jarvis et al. (2014). This tree has a clade **VI+VII** as the earliest diverged Neoaves, the Hoatzin groups with a large clade including **II**+**III**+*S+CR*, and owls are sister to *EHV*. We call to the relationships in this tree the *true model relationships*. We found the overall level of ILS to be relatively high (~58 ARFD between the simulated gene trees and the model species tree). GTEE also appeared to be high, showing ~68 unweighted RF distances between the simulated gene trees and the ML estimates of simulated gene trees, which implies low phylogenetic signal per gene for reconstructing this rapid Neoavian radiation. Even though, coalescent species-trees reconstructed with ASTRAL III using simulated data did recover > 90% of the true model relationships across the tree, the concatenation analyses only recovered ~85% of the true model relationships (Table S4). However, the positions of the Hoatzin and the earliest diverging Neoavian lineage differed from the true model topologies 61% and 44% of the time, respectively, possibly due to the high GTEE and ILS.

The most common topologies for each focal edge in the ML estimates of simulated gene trees were likely to have a single species/lineage sister to the remaining clades (Fig. S7). We conducted EBA for 184 bipartition constraints (see Methods 2.5) using the simulated alignments. Among the bipartitions, there were 63 (earliest Neoaves), 45 (Hoatzin sister), and 14 (owl sister) conflicting hypotheses for each of focal edge (Table 3). The best supported hypotheses usually differed from the true model relationships (Table 3). For example, for the earliest diverging Neoaves, 25 and 28 hypotheses obtained some support from the two sets of simulated exons, respectively. Among them, the model topology (**VI+VII** as earliest Neoaves) is only supported by one or five genes and ranked as the 14th or 18th among the resolutions. The situation was similar for simulated introns and UCEs (Table 3) and for the other two focal edges: the model topology ranks 10th – 17th among the resolutions of the position of the Hoatzin and it ranks second for the position of owls (Table 3) across all data types.

**Table 3.** Simulation results showing the best supported edge

| Simulations | earliest Neoaves<sup>1</sup> (63<sup>#</sup>) | TR | Hoatzin Sister<sup>2</sup> (45) | TR | Owls Sister<sup>3</sup> (14) | TR | **I** (33) | **II** (4) | **III** (16) | **V** (27) |
| --- | --- | --- | --- | --- | --- | --- | --- | --- | --- | --- |
| exon1 | Doves + Turacos | 14 | Sunbittern + Turacos | 12 | Owls + Mousebirds | 2 | **5432\*** | **19746** | **8497** | **5581** |
| exon2 | Cuckoos + Sandgrouse | 18 | Sunbittern + Turacos | 17 | Owls + Mousebirds | 2 | **6438** | **15763** | **7757** | **5416** |
| intron1 | Cuckoos + Doves | 8 | Sunbittern + Turacos | 10 | Owls + Mousebirds | 2 | **4639** | **21835** | **7672** | **6130** |
| intron2 | Cuckoos + Sandgrouse | 14 | Sunbittern + Turacos | 13 | Owls + Mousebirds | 2 | **3805** | **14934** | **7122** | **6537** |
| UCE1 | Cuckoos + Sandgrouse | 14 | Cranes + Flamingos + Grebes | 13 | Owls + Mousebirds | 2 | **4329** | **18031** | **7621** | **5733** |
| UCE2 | Doves + Turacos | 9 | Hoatzin + Nightjars | 17 | Owls + Mousebirds | 2 | **5906** | **20108** | **7616** | **6508** |

NOTE.— TR: the ranking of the model topology of each focal edge in the edge-based analyses. <sup>#</sup>: The number in the parenthesis represents the total number of conflicting constraints for each testing edge. <sup>1</sup>: showing the best hypothesis that is conflict with (VI + VII) + other Neoaves. <sup>2</sup>: showing the best hypothesis that is conflict with Hoatzin + II + III + S + CR. <sup>3</sup>: showing the best hypothesis that is conflict with Owls + EHV. Control edges include I, II, III, and V, which were all best supported by the edge-based analyses. \*Bolded numbers: The difference of $\sum \Delta \ln L$ between the best hypotheses (i.e., I, II, III, and V respectively) and conflicting second best hypotheses for each edge. These large numbers indicate that the control edges can all be recovered with strong support. Clade keys can be found in Table 1 and Figure 3.

In contrast to the focal edges, which were chosen because they exhibited extremely short internodes and dramatic gene tree conflicts, we found that less contentious edges that are united by relatively long internodes (i.e., those with a low level of ILS) received high support across the simulations regardless of data type (Table 3). Examples of these cases included clades **I**, **II**, **III**, and **V** (Fig. 3A), which have been recovered in many previous studies. These clades indicate that EBA can recover the correct relationships when there is sufficient phylogenetic signal and limited ILS. However, the ML estimates of the simulated gene trees can exhibit a surprisingly high degree of disagreement with some strongly corroborated clades. For example, ~20% of the ML estimates of the simulated gene trees failed to recover the (Palaeognathae + Galloanserae) | Neoaves bipartition (i.e., some lineages within Neoaves were sister to Galloanserae). This may seem surprising since the coalescent branch length leading to Neoaves on the model species tree is relatively long (2.03 coalescent units), but we confirmed this level of conflict with the species tree is expected given the multispecies coalescent by examining the simulated gene trees. However, we also note that the amount of ILS in the simulations depends on the coalescent branch lengths estimated by Jarvis et al. (2014). Coalescent branch length calculation is challenging (Forthman et al. 2022) and it is possible that Jarvis et al. (2014) underestimated the coalescent branch lengths; In fact, there is evidence from indels (Houde et al. 2020) that the true coalescent branch length is longer (2.57 coalescent units). Thus, our results represent the behavior of EBA for the case where most incongruence among estimated gene trees reflects ILS rather than GTEE. Nevertheless, the simulations provide guidance regarding the types of errors we might expect for EBA.

### 3.7 Potential limitations of edge-based analyses (EBA)

EBA can be used to assess the relative support for distinct hypotheses, but it does have some limitations. Specifically, edge-based tests are difficult to perform if one wishes to examine a large number of edges (and/or hypotheses regarding the selected edges) due to the high computational load of this approach. This computational burden reflects the need to conduct preliminary tree searches to define the most plausible hypotheses as well as the constrained tree searches. We note that EBA can be viewed as extending another edge-based test, the SH-aLRT (approximate likelihood-ratio test with the nonparametric SH correction, Guindon et al. 2010). The SH-aLRT is used to assess support for edges by comparing the likelihood of the optimal tree with the two alternative trees produced by NNI (Table 1) moves. EBA extends this idea from NNIs to alternatives that: 1) have *a priori* support; or 2) are relatively common in the set of gene trees generated for phylogenomic datasets. Although this makes EBA more flexible it is difficult to devise approximations, like limiting re-optimization to the five edge lengths closest to the focal edge in the optimal tree (which is done in the SH-aLRT).

Another limitation of EBA is the fact the likelihood component of the analyses (Δln*L*) still relies on the substitution model selected for each gene. Obviously, model misspecification can still have an impact on tree topology. In this study, we used the most complex model – time-reversible model (GTR) – to overcome GTEE due to overly simplistic models, but it remains possible that the GTR model is itself inadequate, at least for some loci, so even more complex models should be considered in the future. Another concern in revealed by our simulations is a potential bias toward resolutions of edges that have a single species or lineage sister to the focal taxon, although our simulations are likely to represent a case where ILS is overestimated and GTEE is underestimated. The major benefit of EBA is that it provides information about the support for alternatives to the best-corroborated resolution for each focal edge. Overall, these considerations suggest that EBA might be most appropriate for questions where the number of difficult nodes is limited, when the sets of plausible alternative hypotheses are small, and when one wishes to understand the relative support for each of those plausible alternatives.

## 4. Conclusions

The categorical EBA conducted herein provide a way to examine differences in signal associated with subsets of the data and assess the best-supported resolution of specific nodes. The best-corroborated resolutions of our focal nodes were: 1) owl + *M*; 2) Hoatzin + *S;* and 3) doves are the earliest diverging Neoaves. The best-corroborated hypothesis for owl and Hoatzin was much better supported than the alternatives for all three data types (Fig. 4), although the EBA still revealed substantial conflict. The conflicts evident in the EBA were more complex than the conflicts among analyses of different data types in Jarvis et al. (2014), Reddy et al. (2017), and Braun and Kimball (2021). First, the owls + *EHV* conflict reflects the interaction of GC_CV_ and data type, rather than data type *per se.* Second, the position of Hoatzin was highly variable but there was a data type effect that was not evident in previous studies. Specifically, support for Hoatzin + **V** was strongest in introns, intermediate in UCEs, and weakest for exons. Finally, data type effects were evident for the earliest diverging Neoaves, as expected based on Reddy et al. (2017) and Braun and Kimball (2021), but outlier genes also appeared to play an important role in this part of the tree. There was not a single factor (e.g., GC_CV_ or data type) that played an important role in the resolution of all three focal nodes. Thus, simple solutions like focusing on a specific data type or eliminating genes with high GCCV are unlikely to be a panacea for analyses of avian phylogeny (or, by extension, other nodes in the Tree of Life that have proven to be recalcitrant). We believe that our findings will facilitate further exploration of phylogenetic conflicts using an edge-by-edge approach.

Perhaps one of the most surprising results of this study is the fact that the results of the EBA for the earliest divergence within Neoaves contradict monophyly for two of the magnificent seven clades, which are the superordinal groups that have been consistently identified on the Neoavian tree by recent large-scale studies (Reddy et al. 2017). The consistent recovery of the magnificent seven has led to the view that most or all of these clades are present in the true avian species tree (Suh 2016; Braun et al. 2019; Sangster et al. 2022). However, EBA indicate that at least some of those groups might have less support than we currently believe. For example, signal placing either cuckoos or doves sister to all other Neoaves is stronger than many other alternatives (Figs. 4 and 5). However, the cuckoos sister signal contradicts magnificent seven clade **IV** and the doves sister signal contradicts clade **VI**. There are several interpretations of this result. First, clades **IV** and **VI** could both be present in the true tree, but the conflicting signals evident in EBA reflect the fact that the branches uniting clades **V** and **VI** are especially short and, therefore, “noisy” (e.g., due to ILS). Arguably, this is the simplest hypothesis given that essentially all previous studies (e.g., Jarvis et al. 2014; Prum et al. 2015; Braun and Kimball 2021; Kuhl et al. 2021) agree that the relevant edges are quite short. Second, the fact that signals contradicting those magnificent seven clades are stronger than signals consistent with those clades could provide evidence that clades **IV** and **VI** are present but there is something unusual about the branches uniting those clades. In fact, Houde et al. (2020) examined indel data and they reported that the proportion of indels that conflict with the branch uniting clade **VI** was higher than expected given the relevant edge length in the Jarvis et al. (2014) time tree; they interpreted this as evidence for a transient increase in the effective population size for the common ancestor of that clade. Finally, EBA can be viewed as evidence that the true bird tree lacks either clade **IV** (if the signal supporting the “cuckoos sister” hypothesis is correct) or clade **VI** (if the signal supporting the “doves sister” hypothesis is correct). Indeed, Braun and Kimball (2021) explicitly questioned clade **IV** and Sangster et al. (2022) excluded clade **IV** from the set of strongly corroborated clades (instead, Sangster et al. [2022] named a subset of clade **IV**). These EBA further emphasize the uncertainty regarding clades **IV** and **VI**. Regardless of the best interpretation of the EBA, they provide a way to highlight especially problematic branches that warrant further examination in future phylogenomic studies.

Given the evidence for conflict evident in our EBA we are left with a fundamental question: what is the best-way forward for resolving the nodes in the bird tree that remain uncertain? Obviously, collecting even more data will be important and efforts like the B10K (Zhang 2015; Stiller and Zhang 2019) and the OpenWings (Kimball et al. 2019) projects will provide these larger datasets for birds. However, there is ample evidence that simply expanding the size of phylogenomic datasets will not solve the problem of recalcitrant nodes in avian phylogeny (or the difficult nodes in other parts of the tree of life). Rare genomic changes (Rokas and Holland 2000) may provide an alternative data type that sidesteps the complexities evident in the analyses we present here. In fact, rare genomic changes can avoid (or at least reduce) problems like intralocus recombination, natural selection, and GTEE (Springer and Gatesy 2019). The availability of genome-scale datasets will only increase availability of rare genomic change data and allow the community to move beyond retroposon insertions, which were used even before the phylogenomic era (e.g., Suh et al. 2011), to types of rare genomic changes like numts (Liang et al. 2018) and microinversions (Braun et al. 2011) that have not been used in many phylogenetic studies. These types of data, which are very different from nucleotide sequence data, have the potential to corroborate (or falsify) the hypothesis that the most common resolution in EBA represents a correct resolution of the species tree (at least for some data types). Regardless of whether the hypothesis with the greatest support (whether judged using ∑Δln*L* and ∑Ng) is correct, EBA reveal substantial conflicts within the data. Understanding those conflicts is an important problem in evolutionary biology even if we do not know the true resolution of the tree. We can learn even more if the information about conflicts among genes, like those revealed by our EBA, can be combined with an accurate estimate of the species tree.

## Supplementary material

Supplementary figures and tables can be found online.

## Acknowledgements

We would like to thank Benjamin Winger, Joseph Walker, Nathanael Walker-Hale and Diego Alvarado-Serrano for their constructive comments on the earliest version of the paper. We appreciate other comments from members of the Smith lab and valuable comments from the two anonymous reviewers. This work is supported by grants from the US National Science Foundation (DBI-1458466 and DEB-1207915 to S.A.S., DEB-1241066 and DEB-1146423 to J.C., and DEB-1655683 to E.L.B.), the National Natural Science Foundation of China (No. 32160114 to B.L. and No. 32170460 to N.W.), and by the Frank M. Chapman Memorial Fund of the American Museum of Natural History. N.W. is also supported by Young science and technology talents cultivation project of Inner Mongolia University (No. 21221505).

## Author contributions

Study conception and design: N.W., S.A.S.; Acquisition of data: N.W.; Analysis and interpretation of data: N.W., S.A.S., E.L.B., J.C., B.L.; Drafting of manuscript: N.W., E.L.B., J.C., S.A.S.; Image illustration: B.L., N.W., E.L.B., J.C.

