## Supplementary Figures for "Categorical edge-based analyses of phylogenomic data reveal conflicting signals for difficult relationships in the avian tree"

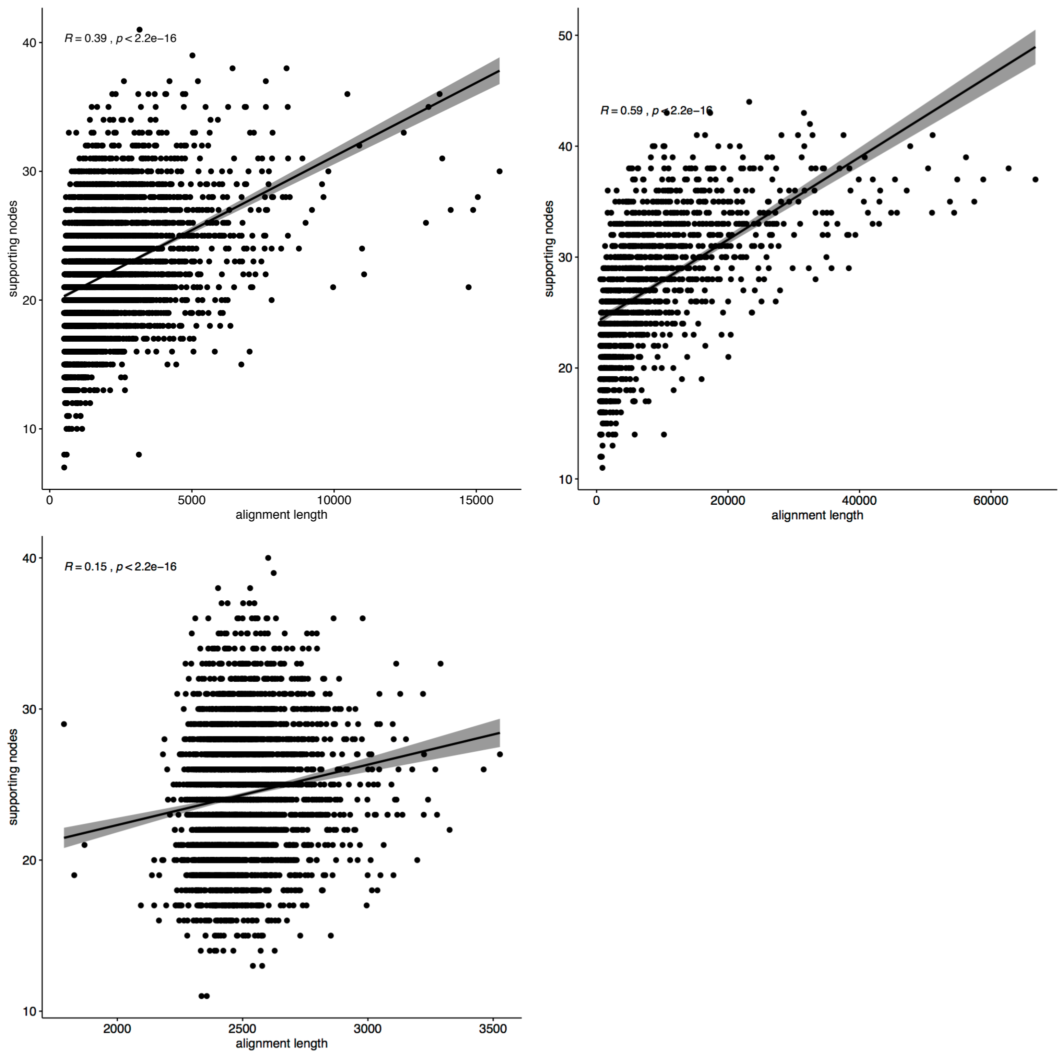


Figure S1. the correlation between supporting node numbers and the alignment length for each dataset (Exons, Introns [upper panel, left to right] and UCEs [lower panel]) in Jarvis et al. (2014).

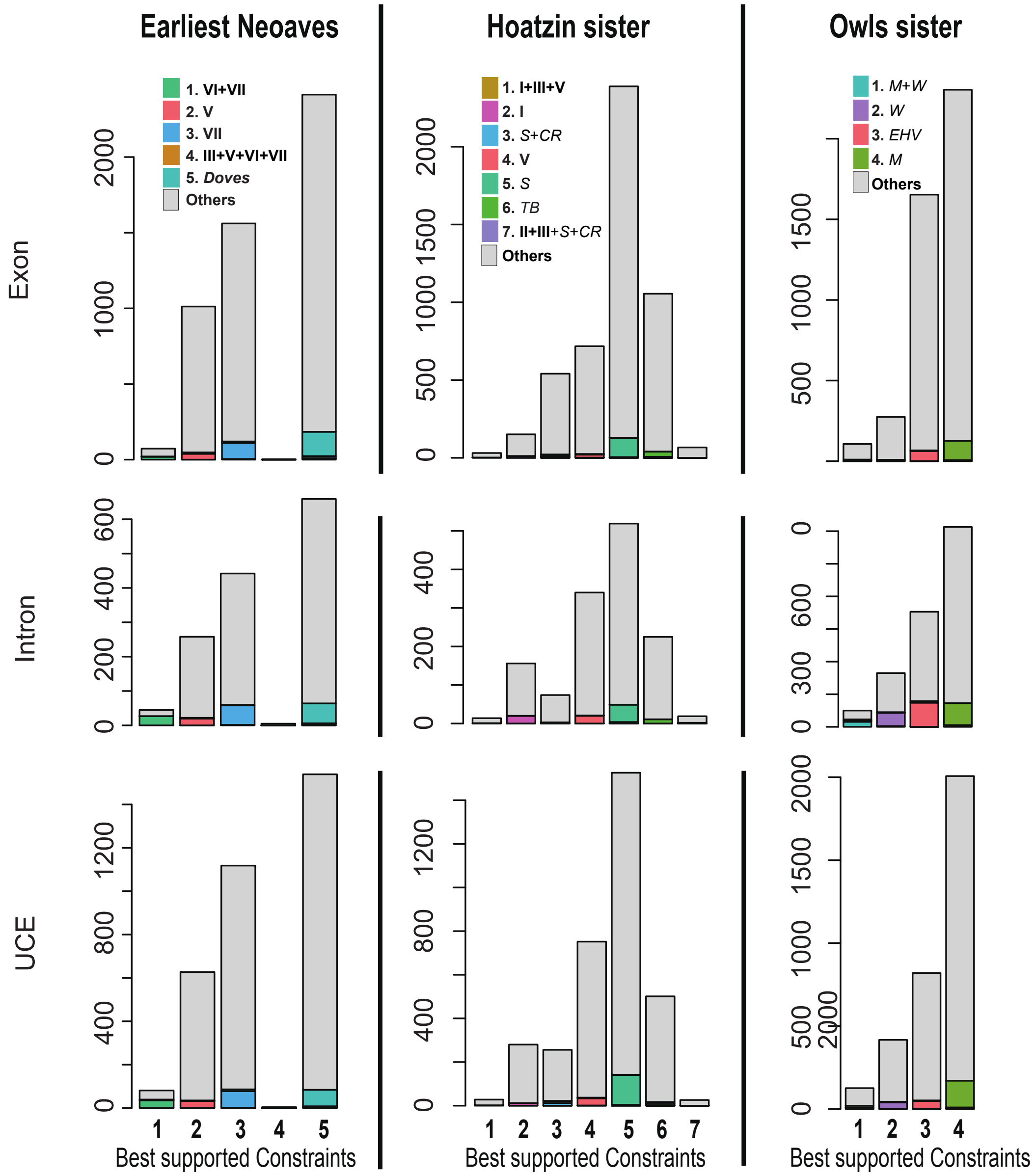


Figure S2: The consistency of the ML gene topology with its best supported constraint. The most genes presented a different ML topology for the focal edge other than its best supporting constraints or any other testing constraints. The Greek numbers (bold) correspond to the "magnificent seven" of Reddy et al. (2017), including **I**: "core" Landbirds; **II**: "core" Waterbirds; **III**: Sunbittern and Tropicbirds; **IV**: Cuckoos, Bustards, and Turacos; **V**: Nightjars, Swifts, Hummingbirds, and allies; **VI**: Doves, Mesites, and Sandgrouse; **VII**: Flamingos and Grebes. Other abbreviations including *S*: Shorebirds; *CR*: Cranes and Rails; *TB*: Turacos and Bustards; *M*: Mousebirds; *W*: Woodpeckers and allies; *EHV*: Eagles, Hawks, and New World Vultures.

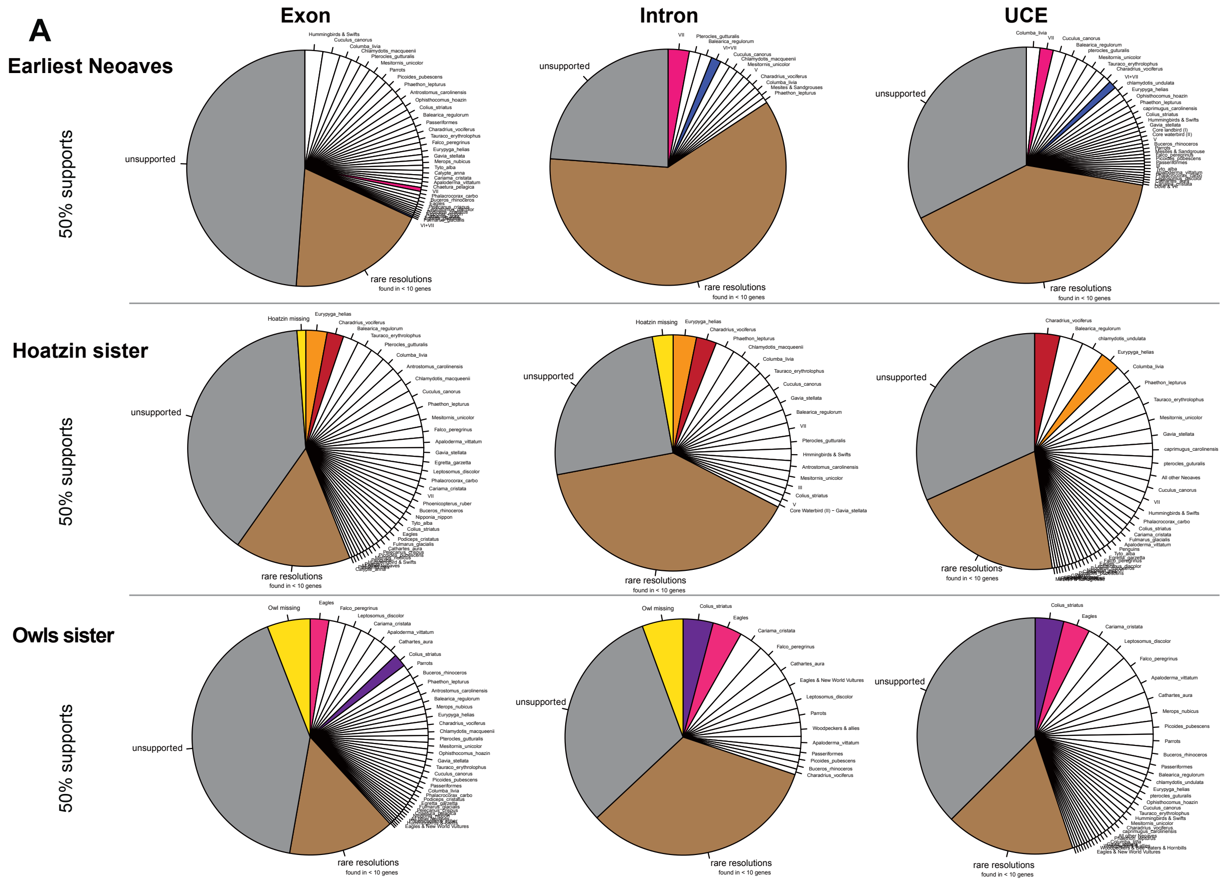


Figure S3. A, Topology distribution of the ML gene trees for the three focal edges using Jarvis et al. (2014) data. unsupported: relationships shown less than 50% UFBS (ultrafast bootstrap support, we use lower support cutoff to show potential topology distributions). Situations are even worse for 95% UFBS cutoff (not shown). rare resolutions: relationships shown in less than 10 gene trees. Hoatzin missing or Owl missing: Hoatzin or Owl is missing in the gene trees.

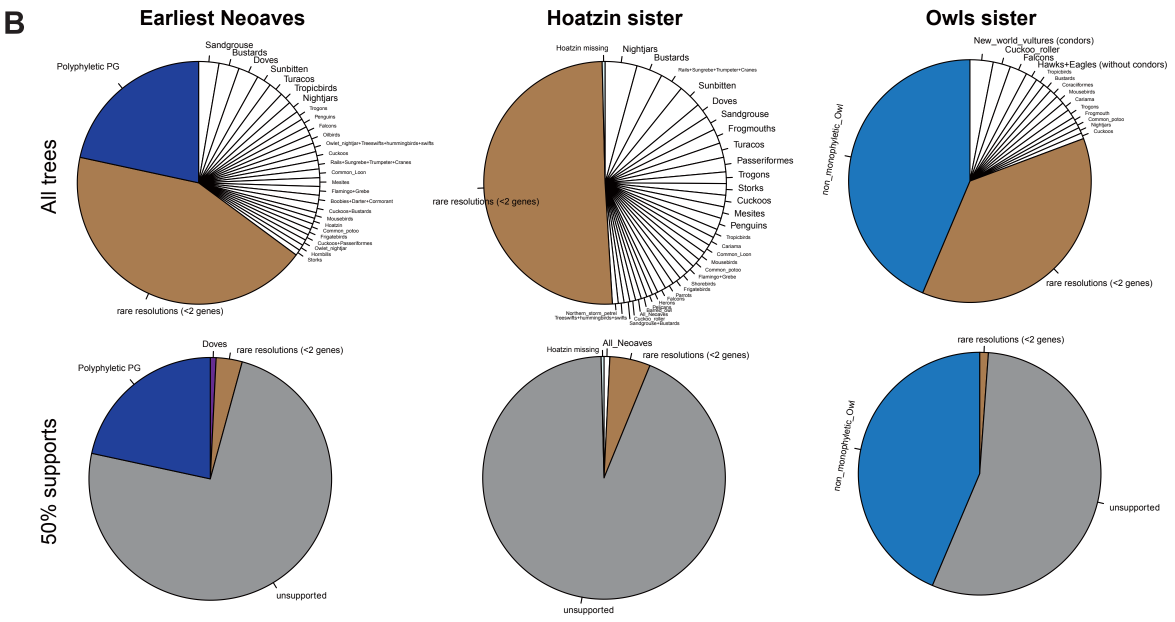


Figure S3. –continued. B, Topology distribution of the ML gene trees for the three focal edges using Prum et al. (2015) data. Upper panel: without constraint bootstrap support. Polyphyletic PG: species within Palaeognathae or Galloanserae are not monophyletic. non_monophyletic_Owl: the two species of Owls used in Prum et al. (2015) are not monophyletic in some gene trees. Lower panel: under 50% bootstrap cutoff. rare resolutions: relationships shown in less than 2 gene trees.

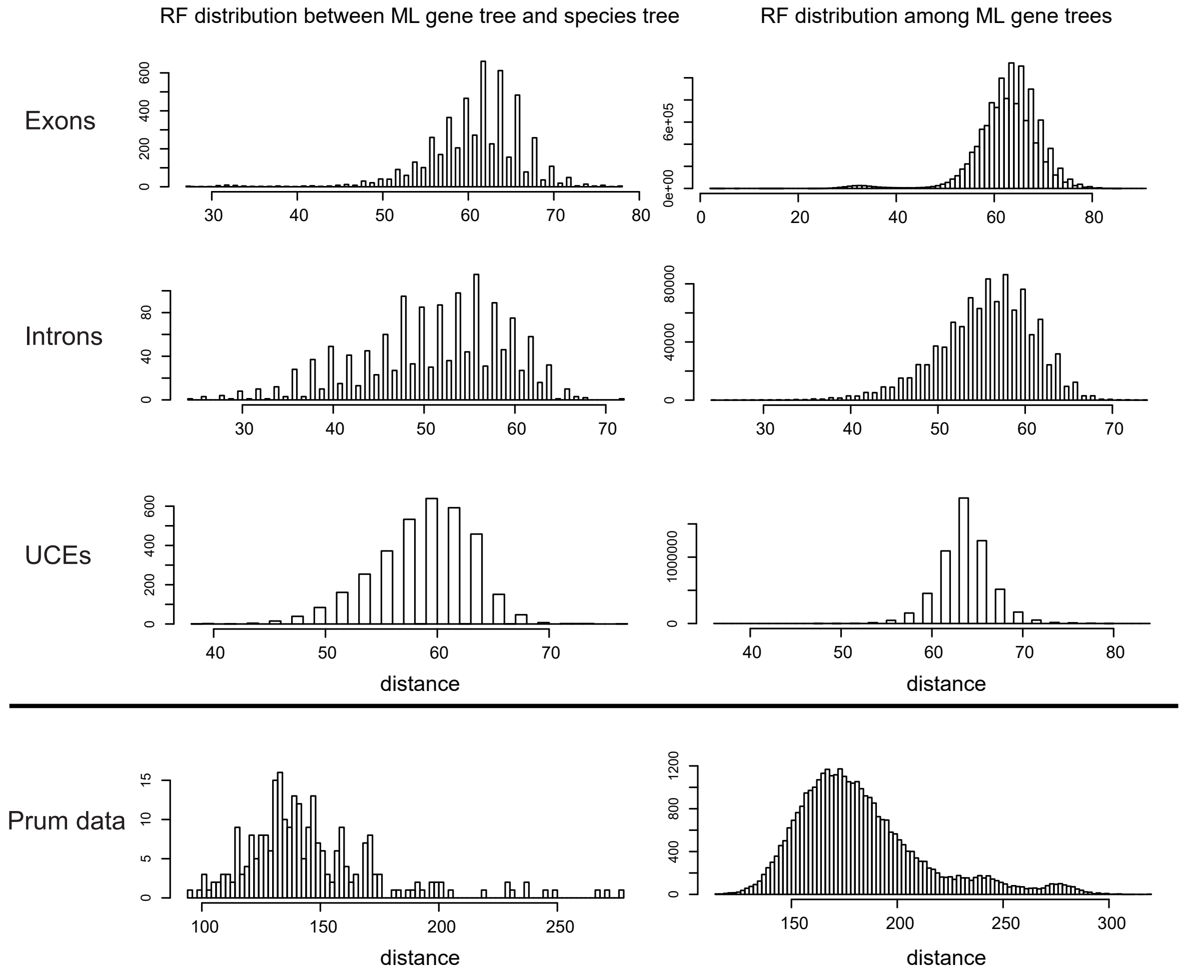


Figure S4. RF (Robinson–Foulds) distances between ML gene trees and species tree (left panel) or among ML gene trees (right panel). For Jarvis et al. (2014) Exons, Introns and UCEs, we use the concatenated TENT tree as species tree. For Prum et al. (2015) data, we use the concatenated RAxML tree as species tree.

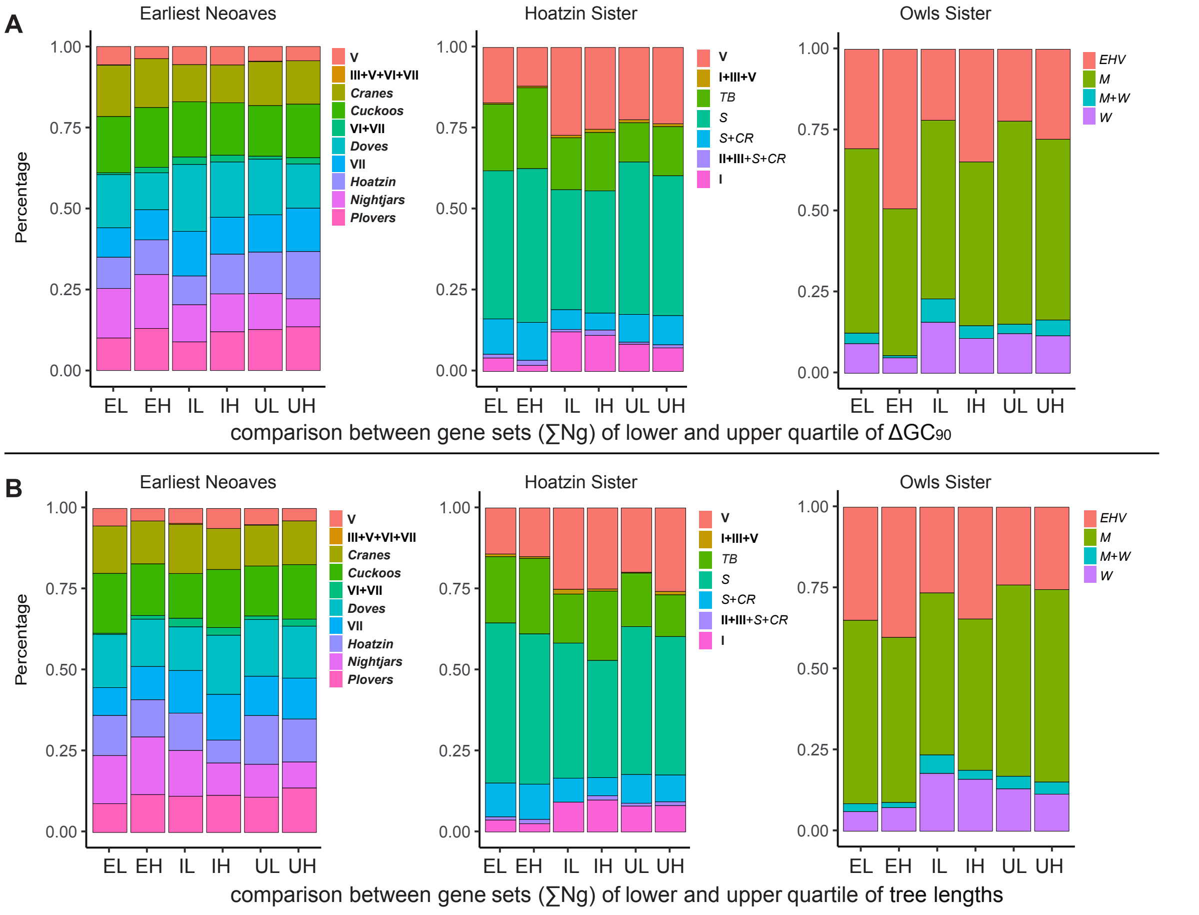


Figure S5. Support comparison among different hypotheses using $\sum$Ng (number of genes) between gene sets of the lower and upper quartiles of A: $\Delta GC_{90}$ (the GC differences between taxa with high [the 95 percentile] and low [the 5 percentile] GC content) or B: tree lengths, for the three focal edges. (EL and EH: exons with lower or higher $\Delta GC_{90}$/tree length; IL and IH: introns with lower or higher $\Delta GC_{90}$/tree length; UL and UH: UCEs with lower or higher $\Delta GC_{90}$/tree length. hypotheses abbreviations are the same as in Figure S2.

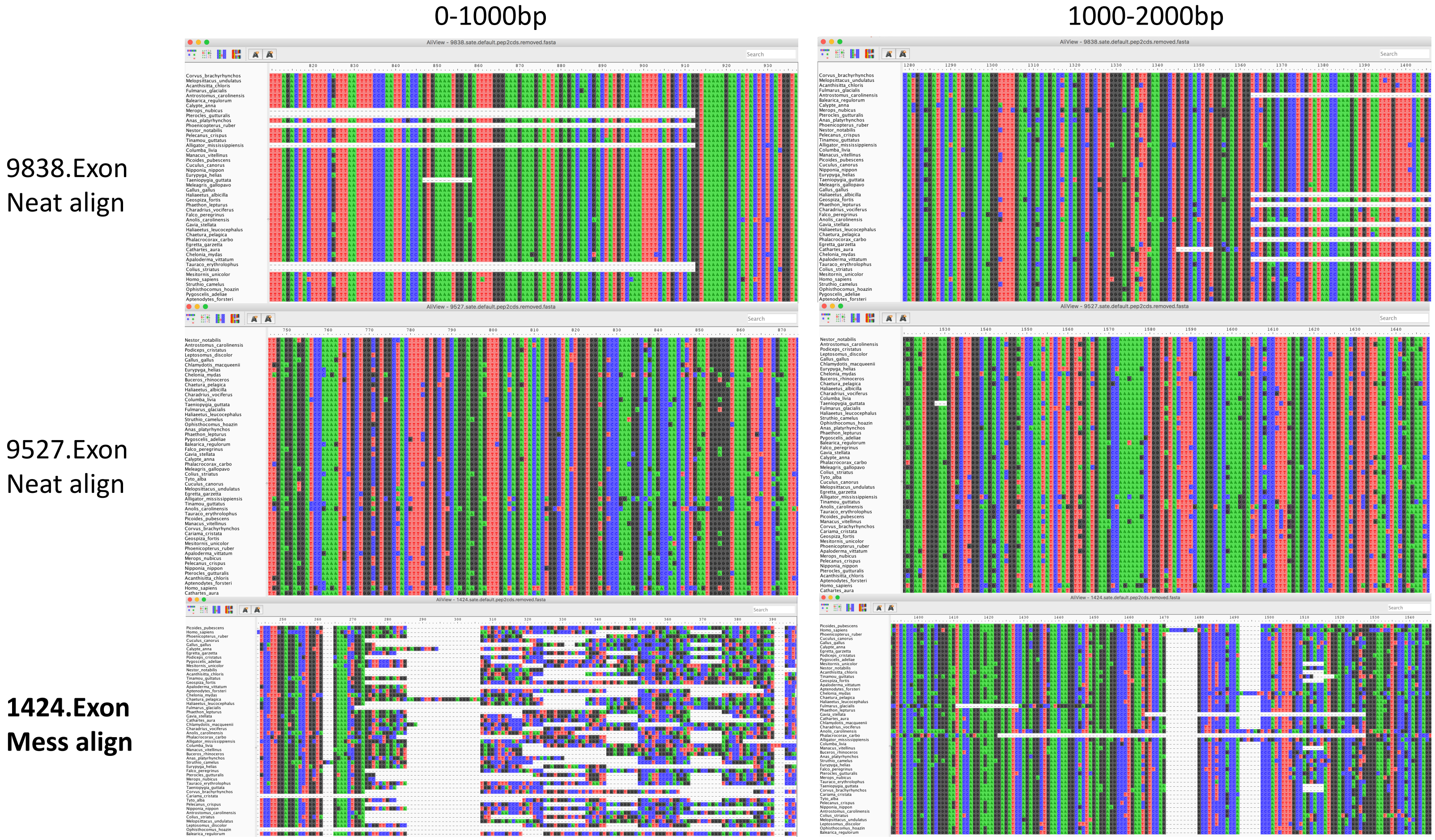


Figure S6. Intragenic alignment comparisons from three exons with intragenic gene tree conflicts, we can see exon locus 1424 have messy alignment from 0-1000bp and the conflicts between the two segments might be corresponded to misalignment.

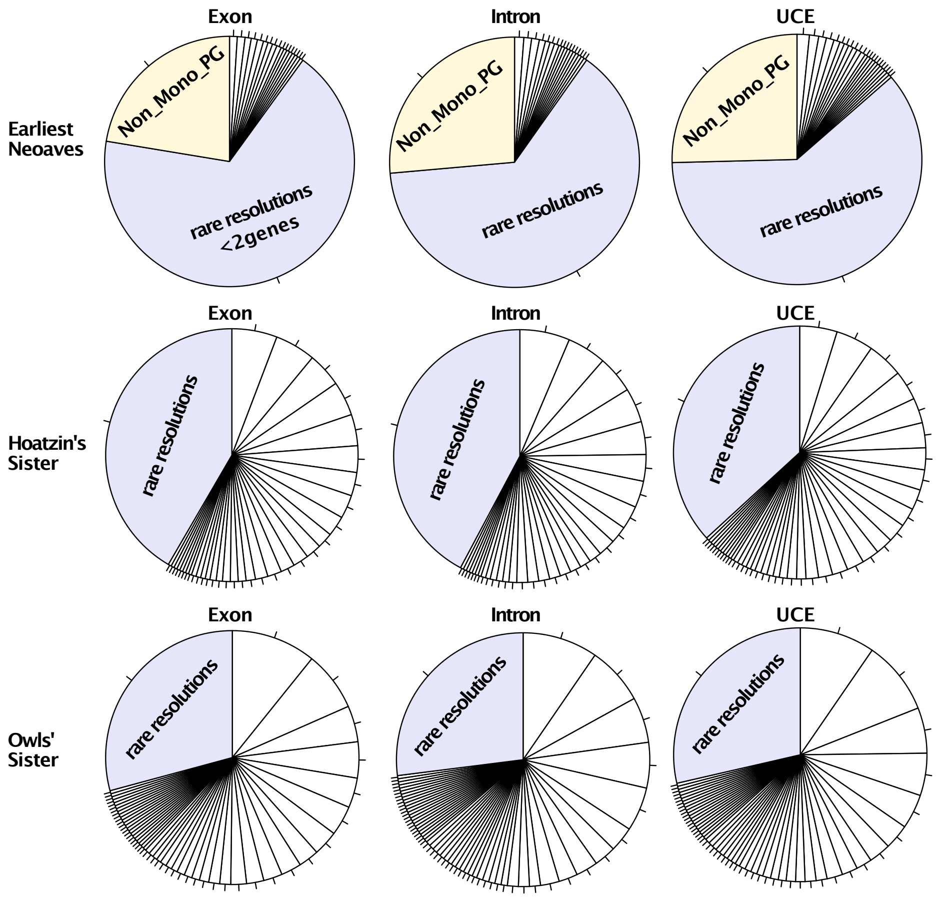


| Exons | No. | Introns | No. | UCEs | No. |
|---|---|---|---|---|---|
| **Earliest Neoaves** | | | | | |
| Exons | No. | Introns | No. | UCEs | No. |
| VII | 5 | VII | 6 | Sandgrouse | 8 |
| Cuckoos | 5 | Bustards | 5 | Doves | 6 |
| Bustards | 4 | Doves | 5 | Bustards | 5 |
| Hoatzin | 4 | Sandgrouse | 4 | VII | 5 |
| Doves | 4 | V | 3 | Killdeer | 4 |
| Grebes | 3 | Cuckoos | 3 | Mesites | 3 |
| Mesites | 3 | Sunbittern | 3 | III+ Doves | 3 |
| Hummingbirds+Swifts | 3 | Nightjars | 2 | Turacos | 3 |
| Sunbittern | 3 | Hoatzin | 2 | Hoatzin | 3 |
| Hoatzin + Killdeer | 2 | Hummingbirds+Swifts | 2 | V | 3 |
| rare resolutions | 338 | rare resolutions | 319 | rare resolutions | 305 |
| **Hoazin's sister** | | | | | |
| Exons | No. | Introns | No. | UCEs | No. |
| Killdeer (shorebirds) | 29 | Killdeer | 32 | Cranes | 24 |
| Sunbittern | 26 | Sunbittern | 25 | Cuckoos | 24 |
| Tropicbirds | 22 | Cranes | 24 | Sunbittern | 23 |
| Cranes | 22 | Tropicbirds | 22 | Killdeer | 19 |
| Turacos | 20 | Doves | 21 | Doves | 16 |
| Bustards | 17 | Cuckoos | 17 | Turacos | 16 |
| Cuckoos | 15 | Turacos | 17 | Tropicbirds | 14 |
| Doves | 15 | Bustards | 15 | Nightjars | 13 |
| Loons | 13 | VII | 11 | Mesites | 13 |
| Nightjars | 10 | Mesites | 11 | III | 12 |
| rare resolutions | 208 | rare resolutions | 211 | rare resolutions | 184 |
| **Owls' sister** | | | | | |
| Exons | No. | Introns | No. | UCEs | No. |
| NW vultures | 54 | NW vultures | 48 | Eagles | 48 |
| Eagles | 38 | Eagles | 37 | NW vultures | 47 |
| Mousebirds | 23 | Seriemas | 29 | Mousebirds | 29 |
| Seriemas | 23 | EHV | 29 | EHV | 27 |
| Cuckoo-roller | 19 | Mousebirds | 28 | Seriemas | 22 |
| EHV | 19 | Falcons | 16 | Cuckoo-roller | 16 |
| Falcons | 16 | Cuckoo-roller | 14 | Falcons | 14 |
| Trogons | 13 | Trogons | 14 | Trogons | 10 |
| Parrots | 13 | Hornbills | 12 | Passerines | 10 |
| Hornbills | 12 | Woodpeckers | 12 | Parrots | 10 |
| rare resolutions | 146 | rare resolutions | 135 | rare resolutions | 143 |

Figure S7. Topology distribution of the *ML estimates of simulated gene trees* for the three focal edges using simulated data. We relax support cutoff to show potential topology distributions. rare resolutions: relationships shown in less than 2 gene trees. Non_Mono_PG: non-monophyletic Palaeognathae and Galloanserae. The right panel showed the top 10 resolutions for each focal edge based on each simulated data type.
