## Supplementary Tables for "Categorical edge-based analyses of phylogenomic data reveal conflicting signals for difficult relationships in the avian tree"

Table S1. Gene filter processes for Jarvis et al. 2014

| Jarvis et al. (2014) | All data | First filter | Second filter | complete matrix | Earliest Neoaves* | Hoatzin* | Owls* |
|---|---|---|---|---|---|---|---|
| Exons | 8295 loci | 7371 loci | 5062 loci | 1300 loci | 5019 loci | 4822 loci | 4194 loci |
| Introns | 2516 loci | 2270 loci | 1409 loci | 185 loci | 1274 loci | 1137 loci | 1134 loci |
| UCEs | 3679 loci | 3679 loci | 3370 loci | 3370 loci | 3197 loci | 3165 loci | 3295 loci |

NOTE.— First filter: remove genes having sequence length < 500bp and large missing taxa (< 42 species after deleting sequences with > 70% gaps in the alignment). Second filter: remove branch length <= 1 expected substitutions per site; remove genes having non-bird species nested within birds, and species from Palaeognathae and Galloanserae nested within Neoaves. * genes filtered based on the compatibility between testing hypotheses and the ML gene trees under 95% UFBS cutoff.

Table S2. Edge-based analyses with Prum et al. (2015) data

| earliest Neoaves | ∑Ng | ∑sigNg | ∑ΔlnL | ∑sigΔlnL |
|---|---|---|---|---|
| **V** | 30 | 18 | 166.31 | 154.13 |
| **VII** | 98 | 82 | **1027.04** | **1011.79** |
| **Doves** | **103** | **85** | 791.67 | 774.35 |
| Hoatzin's sister | ∑Ng | ∑sigNg | ∑ΔlnL | ∑sigΔlnL |
| **I** | 27 | 22 | 173.24 | 169.42 |
| *S* | **182** | **166** | **2972.81** | **2956.90** |
| **V** | 49 | 42 | 1169.73 | 1161.95 |
| Owls' sister | ∑Ng | ∑sigNg | ∑ΔlnL | ∑sigΔlnL |
| *M+W* | 17 | 16 | 278.86 | 278.83 |
| *M* | **142** | **126** | **1588.74** | **1570.19** |
| *EHV* | 99 | 78 | 731.06 | 709.80 |

NOTE.— Hypotheses can be referred to table 2. The first hypothesis in each testing edge is the relationship shown in Prum et al. (2015).
∑Δln*L*: the sum of log likelihood score differences; ∑Ng: the number of genes supporting a specific topology.
∑sigΔln*L*: the sum of log likelihood score differences among genes with significant support (i.e., Δln*L* > 2); ∑sigNg: the number of genes with significant support (i.e., Δln*L* > 2)

Table S3. outlying genes in each testing edge and data types.

| **Exons** | outlying genes | Hypo | Δln*L* | ∑Δln*L* | percent* | BLAST Annotation |
|---|---|---|---|---|---|---|
| earliest Neoaves | 9281.pep2cds | **VII** | 107.3 | 1342.5 | 7.99 | Nipponia nippon BR serine/threonine kinase 2 (BRSK2), transcript variant X1 |
| Owls' sister | 12984.pep2cds | *M+W* | 133.9 | 1046.2 | 12.80 | Cariama cristata myotubularin related protein 6 (MTMR6) |
| | 1672.pep2cds | *M* | 407.2 | 25255.3 | 1.61 | Chelonia mydas vascular cell adhesion molecule 1 (VCAM1) |
| | 599.pep2cds | *M* | 356.5 | 25255.3 | 1.41 | Balearica regulorum gibbericeps WW and C2 domain containing 1 (WWC1) |
| Hoatzin's sister | 9575.pep2cds | **I** | 91.2 | 1111.6 | 8.20 | Charadrius vociferus monocarboxylate transporter 2-like (LOC104285195) |
| | 2661.pep2cds | *S+CR* | 117.0 | 5056.6 | 2.31 | Pygoscelis adeliae trafficking protein, kinesin binding 2 |
| | 10258.pep2cds | *S+CR* | 116.7 | 5056.6 | 2.31 | Colius striatus folliculin interacting protein 2 |
| | 11737.pep2cds | *S+CR* | 103.5 | 5056.6 | 2.05 | Colius striatus KIAA1211 ortholog |
| | 1296.pep2cds | **V** | 110.6 | 4538.7 | 2.44 | Nestor notabilis MICAL-like 2 |
| | 8393.pep2cds | *S* | 308.2 | 23740.9 | 1.30 | Fulmarus glacialis hexokinase 1 (HK1) |
| | 599.pep2cds | *S* | 278.4 | 23740.9 | 1.17 | Balearica regulorum gibbericeps WW and C2 domain containing 1 (WWC1) |
| | 10870.pep2cds | *TB* | 175.7 | 7846.6 | 2.24 | Pelecanus crispus URB2 ribosome biogenesis 2 homolog (S. cerevisiae) (URB2) |
| | 12935.pep2cds | **II+III**+*S+CR* | 192.0 | 953.1 | 20.15 | Fulmarus glacialis lymphocyte cytosolic protein 1 (L-plastin) (LCP1), transcript variant |

| **Introns** | outlying genes | Hypo | Δln*L* | ∑Δln*L* | percent |
|---|---|---|---|---|---|
| earliest Neoaves | 11117.intron | **VI+VII** | 258.7 | 1211.8 | 21.35 |
| | 10376.intron | **VII** | 203.0 | 1651.9 | 12.29 |
| | 11339.intron | Hoazin | 127.1 | 978.9 | 12.98 |
| | 9517.intron | Cranes | 90.1 | 1256.4 | 7.17 |
| | 11084.intron | Cuckoos | 78.0 | 1425.2 | 5.48 |
| Owls' sister | 8210.intron | *EHV* | 128.5 | 2978.3 | 4.31 |
| | 2237.intron | *M* | 186.7 | 6584.0 | 2.84 |
| | 11557.intron | *M* | 185.4 | 6584.0 | 2.82 |
| | 12053.intron | *M* | 154.9 | 6584.0 | 2.35 |
| Hoatzin's sister | 7742.intron | **I** | 291.1 | 4768.3 | 6.10 |
| | 3157.intron | *S+CR* | 450.1 | 1528.1 | 29.45 |
| | 4121.intron | **V** | 286.1 | 6966.8 | 4.11 |
| | 2905.intron | **V** | 216.7 | 6966.8 | 3.11 |

| | | | | | |
|---|---|---|---|---|---|
| | 10828.intron | *S* | 218.1 | 8437.9 | 2.58 |
| | 12765.intron | *TB* | 168.2 | 2346.2 | 7.17 |
| | 12666.intron | **II**+**III**+*S*+*CR* | 210.5 | 403.2 | 52.20 |
| **UCEs** | outlying genes | Hypo | ΔlnL | ∑ΔlnL | percent |
| earliest Neoaves | chr4_13270_s.UCE | **VI**+**VII** | 101.4 | 921.6 | 11.00 |
| | chr1_28895_s.UCE | **V** | 351.0 | 887.3 | 39.56 |
| | chr8_1881_s.UCE | **VII** | 665.7 | 2106.1 | 31.61 |
| Owls' sister | chr12_2272_s.UCE | *W* | 344.1 | 2335.6 | 14.73 |
| | chr1_1378_s.UCE | *EHV* | 445.0 | 5282.7 | 8.42 |
| | chr1_25_s.UCE | *EHV* | 217.8 | 5282.7 | 4.12 |
| | chr1_5460_s.UCE | *EHV* | 188.0 | 5282.7 | 3.56 |
| | chr2_6757_s.UCE | *EHV* | 162.9 | 5282.7 | 3.08 |
| | chr20_1620_s.UCE | *EHV* | 162.7 | 5282.7 | 3.08 |
| | chr11_2183_s.UCE | *EHV* | 125.1 | 5282.7 | 2.37 |
| | chr11_2131_s.UCE | *M* | 476.4 | 15591.5 | 3.06 |
| | chr2_7444_s.UCE | *M* | 264.9 | 15591.5 | 1.70 |
| | chr15_4578_s.UCE | *M* | 166.1 | 15591.5 | 1.07 |
| Hoatzin's sister | chr6_11241_s.UCE | **I** | 510.7 | 3492.6 | 14.62 |
| | chr8_12492_s.UCE | *S*+*CR* | 136.7 | 2543.7 | 5.38 |
| | chr2_7444_s.UCE | *S*+*CR* | 135.0 | 2543.7 | 5.31 |
| | chr6_10764_s.UCE | **V** | 135.1 | 5979.0 | 2.26 |
| | chr1_1131_s.UCE | *S* | 449.3 | 14925.7 | 3.01 |
| | chr14_3795_s.UCE | *S* | 373.8 | 14925.7 | 2.50 |
| | chr11_2355_s.UCE | *S* | 218.0 | 14925.7 | 1.46 |
| | chr2_17651_s.UCE | *TB* | 174.1 | 3671.4 | 4.74 |

NOTE.— Hypotheses including **I**: core Landbirds; **II**: core waterbirds; **III**: Sunbittern + Tropicbirds; **V**: Nightjars + Swifts + Hummingbirds; **VI**: Doves + Sandgrouse + Mesites **VII**: Flamingos + Grebes; *S*: Shorebirds (represented by Plovers); *CR*: Cranes and Rails; *TB*: Turacos + Bustards; *EHV*: Eagles, Hawks, and New World Vultures; *M*: Mousebirds; *W*: Woodpeckers and allies.
*percent means ΔlnL/∑ΔlnL.

Table S4. Edge recovery based on simulation data.

| Simulation analyses | Focal edges | | | The magnificent seven in Reddy et al. (2017) | | | | | | |
|---|---|---|---|---|---|---|---|---|---|---|
| TENT.MP-EST.binned.tre (*model species tree*) | earliest Neoaves | Hoatzin's position | Owls' position | **I** | **II** | **III** | **IV** | **V** | **VI** | **VII** |
| Exon_simAstral1* | ✓ | Hoatzin + **II** + **III** + *CR* | ✓ | ✓ | ✓ | ✓ | ✓ | ✓ | ✓ | ✓ |
| Exon_MLAstral1# | ✓ | Hoatzin + **III** | ✓ | ✓ | ✓ | ✓ | ✓ | ✓ | ✓ | ✓ |
| Exon_concatenation tree1 | ✓ | Hoatzin + *S* + *CR* | ✓ | ✓ | ✓ | ✓ | ✓ | ✓ | ✓ | ✓ |
| Exon_simAstral2 | ✓ | Hoatzin + **I** | ✓ | ✓ | ✓ | ✓ | ✓ | ✓ | ✓ | ✓ |
| Exon_MLAstral2 | **VI** | ✓$ | ✓ | ✓ | ✓ | ✓ | ✓ | ✓ | ✓ | ✓ |
| Exon_concatenation tree2 | **VI** | Hoatzin + **I** + **V** | ✓ | ✓ | ✓ | ✓ | ✓ | ✓ | ✓ | ✓ |
| Intron_simAstral1 | ✓ | Hoatzin + **I** + **II** | ✓ | ✓ | ✓ | ✓ | ✓ | ✓ | ✓ | ✓ |
| Intron_MLAstral1 | ✓ | ✓ | ✓ | ✓ | ✓ | ✓ | ✓ | ✓ | ✓ | ✓ |
| Intron_concatenation tree1 | **VII** | Hoatzin + **I** + **IV + V** | ✓ | ✓ | ✓ | ✓ | ✓ | ✓ | ✓ | ✓ |
| Intron_simAstral2 | ✓ | ✓ | ✓ | ✓ | ✓ | ✓ | ✓ | ✓ | ✓ | ✓ |
| Intron_MLAstral2 | ✓$ | ✓ | ✓ | ✓ | ✓ | ✓ | ✓ | ✓ | X | ✓ |
| Intron_concatenation tree2 | **VII** | Hoatzin + *S* + **V** | ✓ | ✓ | ✓ | ✓ | X | ✓ | X | ✓ |
| UCE_simAstral1 | **VI** | ✓ | ✓ | ✓ | ✓ | ✓ | ✓ | ✓ | ✓ | ✓ |
| UCE_MLAstral1 | **VI** | ✓ | ✓ | ✓ | ✓ | ✓ | ✓ | ✓ | ✓ | ✓ |
| UCE_concatenation tree1 | Sandgrouse + Mesites | Hoatzin + **II** + **III** | ✓ | ✓ | ✓ | ✓ | ✓ | ✓ | X | ✓ |
| UCE_simAstral2 | ✓$ | ✓ | ✓ | ✓ | ✓ | ✓ | ✓ | ✓ | X | ✓ |
| UCE_MLAstral2 | ✓$ | Hoatzin + *S* | ✓ | ✓ | ✓ | ✓ | ✓ | ✓ | X | ✓ |
| UCE_concatenation tree2 | **VII** | Hoatzin + **II** + *CR* | ✓ | ✓ | ✓ | ✓ | X | ✓ | ✓ | ✓ |

NOTE.— *: simAstral: the coalescent species tree with ASTRAL III using the rescaled simulated gene trees generated from the model species tree (i.e., TENT.MP-EST.binned.tre of Jarvis et al. [2014]). #: MLAstral: the coalescent species tree with ASTRAL III using the ML estimates of simulated gene trees based on the simulated alignments. $: the internal relationships within the group are different from model species tree.

concatenation analyses were conducted using the simulated alignments in RAxML 8.2.11.

The magnificent seven are superordinal groups from Reddy et al. 2017, including **I**: core Landbirds; **II**: core Waterbirds; **III**: Tropicbirds + Sunbittern; **IV**: Cuckoos + Turacos + Bustards; **V**: Nightjars + Swifts + Hummingbirds; **VI**: Doves + Sandgrouse + Mesites; **VII**: Flamingos + Grebes.

"✓" means the relationships on the tree match the topologies on the model species tree. Specifically, the model species tree has a clade **VI+VII** as the earliest diverged Neoaves, the Hoatzin groups with a large clade including **II**+**III**+*S* (Shorebirds) +*CR* (Cranes and Rails), and owls are sister to *EHV* (Eagles, Hawks and New World Vultures). X means the superordinal group was not recovered.
